# Biological constraints on stereotaxic targeting of functionally-defined cortical areas

**DOI:** 10.1101/2022.03.04.483047

**Authors:** Divya P Narayanan, Hiroaki Tsukano, Amber M Kline, Koun Onodera, Hiroyuki K Kato

## Abstract

Understanding computational principles in hierarchically organized sensory systems requires functional parcellation of brain structures and their precise targeting for manipulations. Although brain atlases are widely used to infer area locations in the mouse neocortex, it has been unclear whether stereotaxic coordinates based on standardized brain morphology accurately represent functional domains in individual animals. Here, we used intrinsic signal imaging to evaluate the accuracy of area delineation in the atlas by mapping functionally-identified auditory cortices onto bregma-based stereotaxic coordinates. We found that auditory cortices in the brain atlas correlated poorly with the true complexity of functional area boundaries. Inter-animal variability in functional area locations predicted surprisingly high error rates in stereotaxic targeting with atlas coordinates. This variability was not simply attributed to brain sizes or suture irregularities but instead reflected differences in cortical geography across animals. Our data thus indicate that functional mapping in individual animals is essential for dissecting cortical area-specific roles with high precision.

## Introduction

The cerebral cortex consists of numerous functionally-specialized areas that form hierarchically organized streams to support diverse sensory computation (Felleman and Van Essen, 1991; Kaas and Hackett, 2000; Maunsell and Newsome, 1987; Rauschecker and Tian, 2000). One of the central goals in neuroscience is to dissect this complex network and link the neural dynamics of individual cortical areas to perceptual behaviors. To achieve this goal, it is essential to parcellate functionally-defined cortical areas and further conduct area- targeted experiments, such as neural recording (Guo et al., 2012; Joachimsthaler et al., 2014; Siegle et al., 2021), connectivity tracing (Llano and Sherman, 2008; Oh et al., 2014; Schreiner and Winer, 2007; Wang and Burkhalter, 2007), and functional manipulation during behaviors (Ceballo et al., 2019; Kline et al., 2021). Over the last decade, the mouse has emerged as an essential model system to study cortical sensory pathways with the development of genetic and viral tools for circuit dissection (Luo et al., 2018; Roth, 2016; Tervo et al., 2016; Wickersham et al., 2007; Yizhar et al., 2011; Zingg et al., 2017). Although the compact size of mouse brains is beneficial to the comprehensive understanding of sensory processing pathways, it simultaneously necessitates additional precision in targeting small cortical domains for experiments such as the insertion of recording electrodes or virus injection pipettes.

The lissencephalic mouse neocortex lacks structural landmarks to accurately delineate functional area borders. Therefore, the locations of cortical areas are often inferred from standardized brain structures, such as the Paxinos and Franklin Mouse Brain Atlas (Paxinos and Franklin, 2019, 2012) and the Allen Mouse Brain Common Coordinate Framework (Q. Wang et al., 2020). In particular, stereotaxic coordinates with respect to bregma, a suture landmark on the skull, in the Paxinos Atlas have been widely used to target specific cortical regions. In the Paxinos Brain Atlas, cortical area boundaries were estimated based on the immunostaining patterns of neurofilaments and calcium-binding proteins (Cruikshank et al., 2001; Oh et al., 2014; Paxinos and Franklin, 2019, 2012; Wang et al., 2011). With this strategy, mouse brain atlases have traditionally divided the auditory cortex into three large subregions—the primary auditory field (Au1) and two higher-order areas, the dorsal auditory field (AuD) and ventral auditory field (AuV). However, this simple area segmentation contrasts with the complex arrangement of functionally-mapped auditory cortices, which includes at least four tonotopic areas and one or two non-tonotopic areas (Aponte et al., 2021; Guo et al., 2012; Issa et al., 2014; Joachimsthaler et al., 2014; Kline et al., 2021; Liu et al., 2019; Romero et al., 2020; Stiebler et al., 1997; Tsukano et al., 2015, 2017). This discrepancy in area parcellation schemes calls into question how accurately stereotaxic targeting based on the brain atlases captures functionally-defined cortical areas in individual animals.

Furthermore, stereotaxic targeting based on brain atlases can be confounded by inter-individual anatomical differences in both the brain and skull. The brain atlases were developed by generating an average template from a large number of animals (Paxinos Atlas: 26 mice, Allen Brain Atlas: 1,675 mice). This procedure inevitably discards any inter-animal structural variability existing in the population. For example, stereotaxic coordinates measured from bregma can be affected by biological variabilities such as irregular suture patterns (Blasiak et al., 2010; Whishaw et al., 1977; Zhou et al., 2020), brain size differences (Paxinos et al., 1985; Wahlsten et al., 1975; N. Wang et al., 2020), and variability in the relative positioning of functional areas within the cortex (Garrett et al., 2014; Waters et al., 2019). These variabilities limit the accuracy of stereotaxic targeting and may result in misinterpretation of experimental data, especially in small brain areas such as the auditory cortex. However, direct comparison between the brain atlas and functional maps and systematic quantification of the inter-animal variability in the bregma-based coordinates of functional areas have yet to be conducted in any cortical areas.

In this study, using intrinsic signal imaging, a non-invasive functional mapping free from tissue damage-related distortion, we directly measure the stereotaxic locations of functionally-identified auditory cortices in a large group of mice, including multiple strains and both sexes. We demonstrate that the shape and size of functional cortical areas are remarkably variable across individuals. Most strikingly, the stereotaxic location of the auditory cortex shows inter-animal variability as large as one millimeter along both the anteroposterior (AP) and dorsoventral (DV) axes. As a consequence, direct comparison between the brain atlas and functional maps reveals substantial mismatches, which highly limit the accuracy of stereotaxic targeting. Our results indicate the necessity of functional mapping in individual animals instead of relying on stereotaxic coordinates based on standardized brain atlases. To encourage the use of intrinsic signal imaging as a standard mapping method prior to cortical targeting, we provide a detailed protocol, including both the optical setup and surgical procedures. We hope this information will help researchers perform precise areal targeting and accelerate the future dissection of cortical networks underlying perceptual behaviors.

## Results

### The absolute stereotaxic location of functionally-identified mouse auditory cortex varies across animals

To measure the variability in the stereotaxic location of functionally-identified auditory cortex, we compared the cortical map generated by intrinsic signal imaging to the location of commonly used stereotaxic landmarks, bregma and lambda (Paxinos and Franklin, 2019, 2012). We used both sexes (male: n = 25 mice; female: n = 16) as well as multiple wildtype and transgenic strains (C57BL/6J (B6): n = 14; CBA: n = 13; PV-Cre×Ai9: n = 7; VGAT-Cre×Ai9: n = 6) to investigate the influence of genetic backgrounds. Using bregma and lambda as a guide, we marked three stereotaxic reference points ((Posterior, Ventral) = (−2.5, 1.5), (−3.5, 1.5), (−3.5, 2.0); coordinates are in millimeters from bregma unless otherwise specified. See Methods for alignment procedures) near the auditory cortex with black ink on the skull surface (Figure 1A). To compensate for differences in brain size across animals, we normalized the bregma–lambda distance to 4.2 mm, a standard distance for adult B6 males (see the next paragraph for justification) (Paxinos and Franklin, 2019). In the same animals, we performed intrinsic signal imaging through the skull to map cortical areas responsive to pure tones of three frequencies (3, 10, and 30 kHz, 75 dB SPL, 1 s) (Figure 1A, B). Based on the intrinsic signal imaging maps, we performed semiautomated sorting of area boundaries into the primary auditory cortex (A1), ventral auditory field (VAF), secondary auditory cortex (A2), and anterior auditory field (AAF) (Figure 1C; threshold at 60% peak response amplitudes; see Methods). We allowed A1 and VAF to overlap as we observed convergence of these two areas at their low-frequency poles in most animals (Aponte et al., 2021; Issa et al., 2014; Kline et al., 2021; Liu et al., 2019; Romero et al., 2020). Surprisingly, direct comparison between this functional map and the stereotaxic reference points on the skull revealed high variability in the stereotaxic location of functionally-identified auditory cortical areas (Figure 1D). To quantify this variability, the three stereotaxic reference points were used to integrate the functional maps into the stereotaxic coordinate system. When we plotted the centroids of individual frequency domains from all mice onto this coordinate system, we found inter-animal variability as large as 1 mm along both AP and DV axes (Figure 1E, n = 41 mice).

**Figure 1.**
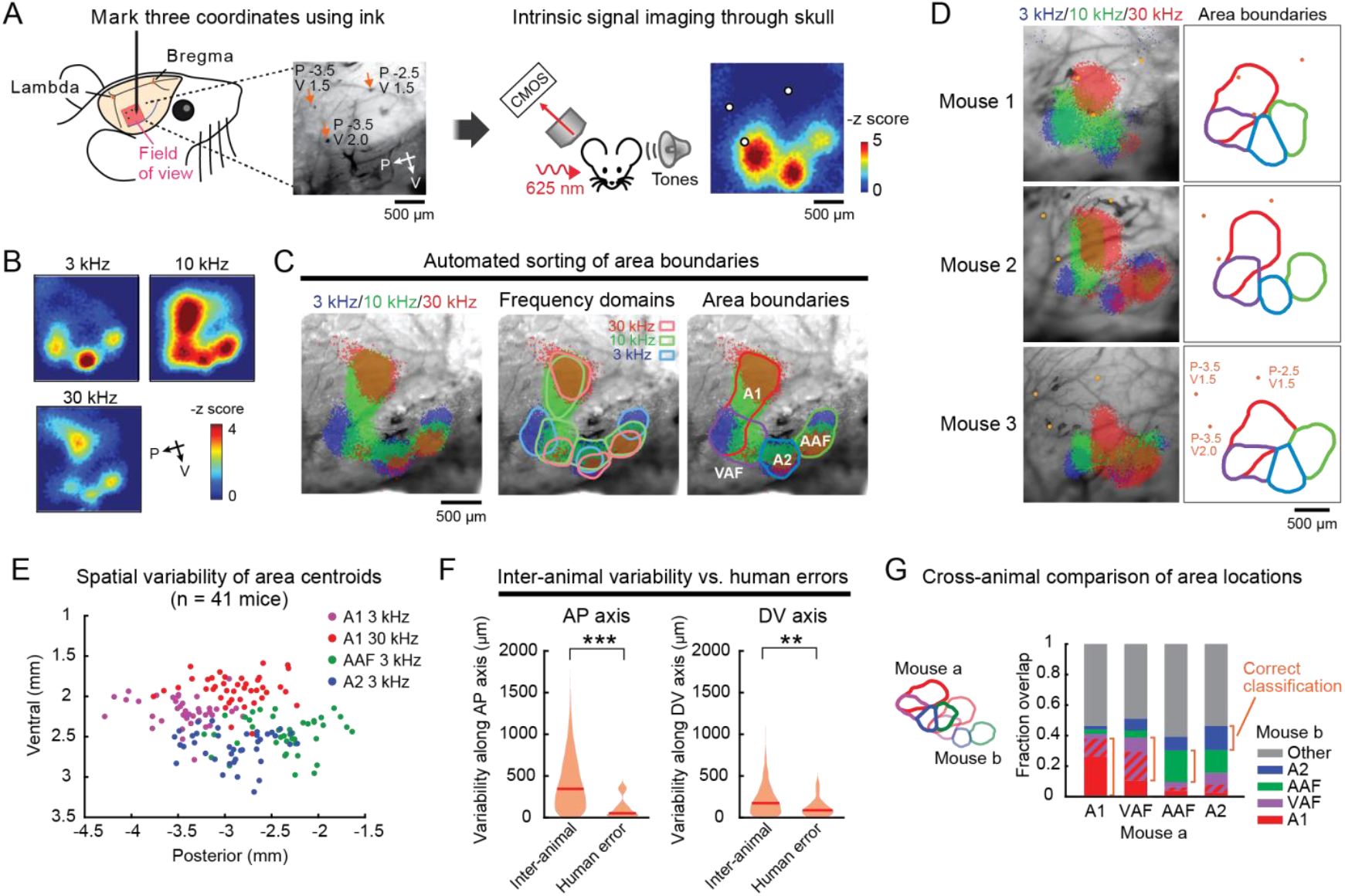
Stereotaxic locations of functionally-identified cortical areas vary across individuals. **(A)** Left, Experimental setup. The stereotaxic coordinates of (P, V) = (−2.5, 1.5), (−3.5, 1.5), (−3.5, 2.0) are marked on the skull surface. Right, intrinsic signal imaging setup. A representative heat map of 3 kHz tone response is overlayed with the extracted mark positions. **(B)** Responses to 3, 10, 30 kHz tones visualized in the right auditory cortex of a representative mouse. Heat maps show z-scored response amplitudes. **(C)** Semiautomated area segmentation in the same mouse as (B). Left, thresholded intrinsic signal image is superimposed on cortical vasculature imaged through the skull. Middle, boundaries of frequency domains for individual cortical areas. Right, area boundaries determined by merging frequency domains. **(D)** Representative functional maps of auditory cortical areas in three mice, showing inter-animal variability in their locations relative to the stereotaxic reference points (orange dots). **(E)** Scatter plot showing the distribution of functionally-identified frequency domain centroids (A1 3 kHz, A1 30 kHz, AAF 3 kHz, and A2 3 kHz) across mice (n = 41 mice). **(F)** Violin plots comparing inter-animal variability and human error along anteroposterior (left) and dorsoventral (right) axes. Inter-animal variability is measured as distances between corresponding frequency domain centroids across mice (n = 3240 centroid pairs). Human error is measured as distances between stereotaxic reference points marked by two experimenters in the same animals (n = 16 mice). Red lines are median. ***p = 3.45×10^−7^, **p = 0.0036, Wilcoxon rank sum test. **(G)** Bar plots displaying cross-animal overlap of functional area locations (n = 41 mice, 1640 mouse pairs). The overlapping region between A1 and VAF is indicated in stripes. Orange brackets show the fraction of correct classification across animals.

Although commonly used for stereotaxic targeting, bregma-based absolute coordinates without size normalization resulted in even larger variability than the size-normalized data (p = 7.15×10^−10^, Wilcoxon rank sum test with Bonferroni correction; Supplementary Figure 1). Lambda-based absolute coordinates gave similar results to the size-normalized data, likely due to the proximity of the auditory cortex to lambda. Scaling DV coordinates based on bregma–lambda distance did not reduce variability. For the rest of the analysis, we chose to normalize only AP coordinates by bregma–lambda distance—the method which resulted in the smallest variability. Therefore, the variability reported in this study is likely an underestimate compared to the standard method of using bregma-based absolute coordinates. When multiple experimenters marked the stereotaxic reference points in the same animals, the inter-experimenter variability of ink locations was significantly smaller than the inter-animal variability of area centroids (Figure 1F, AP axis: inter-animal, 410 ± 5 μm, n = 3240 centroid pairs, inter-experimenter, 106 ± 28 μm, n = 16 mice, p = 3.45×10^−7^; DV axis: inter-animal, 215 ± 3 μm, inter-experimenter, 114 ± 25 μm, p = 0.0036; Wilcoxon rank sum test). This result indicates that human error in identifying stereotaxic landmarks cannot account for the observed variability, suggesting a biological source of variation in auditory cortex stereotaxic locations.

We observed intermixing between centroids of different areas across animals, suggesting the unreliability in transferring the functional area locations from one animal to another (Figure 1E). To quantify how well functional area locations are conserved across mice, we measured the overlap of each functional area mask across pairs of animals. We found that the fraction of overlap between the absolute location of the corresponding auditory cortical areas is surprisingly low across animals (Figure 1G, A1–A1: 38%, VAF– VAF: 26%, AAF–AAF: 21%, A2–A2: 16%). Moreover, the auditory cortex location in one mouse fell outside that of another mouse over 50% of the time. Overall, this marked variability in the stereotaxic location of auditory cortices suggests that the location of functional cortical areas cannot be generalized across individuals.

### Brain atlas coordinates are inaccurate for targeting functional auditory cortical areas

Reference maps of brain structures, such as the Paxinos Brain Atlas (Paxinos and Franklin, 2019, 2012), are commonly used to target the auditory cortex using stereotaxic coordinates. Generally, Au1 has been used to target the primary auditory cortex while AuD and AuV are considered secondary cortices. More specifically, AuV has been considered as a proxy for functional A2. However, the observed inter-animal variability of auditory cortex locations raises the question of how accurately these standardized atlas areas represent the functionally-identified areas in individual animals. Therefore, we next determined how functionally- identified auditory cortical areas map onto three auditory cortex subdivisions in the Paxinos Brain Atlas. We focused on the Paxinos Brain Atlas since the Allen Brain Atlas does not indicate a bregma location. We generated a topographical surface map of atlas-defined cortical areas by first extracting the DV coordinates of area borders from each coronal atlas section and connecting them along the AP axis (Figure 2A, Supplementary Figure 2). When we plotted the probability distribution of functionally-identified A1, VAF, A2, and AAF area masks across mice onto the topographical atlas map, the functionally-generated map was mostly encompassed by the brain atlas auditory subdivisions (Figure 2B; n = 41 mice), validating the overall accuracy of the atlas coordinates in locating the auditory cortex at a population level. Importantly, however, when we looked at individual mice, we observed high variability in how the functional areas map onto the brain atlas (Figure 2C, D). We quantified the relationship between functionally-identified areas (A1, A2, AAF, and VAF) and atlas areas (Au1, AuD, and AuV) by both mapping the functional areas onto atlas areas (Figure 2E) and vice versa (Figure 2F). We observed three notable dissociations of our functional mapping data from the common usage of the brain atlas. First, functionally-identified primary areas A1, VAF, and AAF had large overlaps with AuD and AuV, while secondary region A2 had substantial overlap with Au1 (Figure 2E). As a result, if we rely on the brain atlas, neural events in A1 (such as neural activity or gene expression) would be correctly classified as the primary auditory cortex only about half of the time, and only slightly higher for events in A2 being classified as the secondary auditory cortex (Figure 2E, A1→Au1: 48 ± 4%, A2→AuV: 54 ± 4%). Therefore, the simple binary classification of Au1 as the primary and AuD/AuV as the secondary cortex is misrepresentative. Second, functionally-identified A2 in individual mice is much smaller than AuV, which already suggests that using AuV to target the secondary auditory cortex will result in a high fraction of error (Figure 2F). Indeed, AuV had a larger overlap with the primary area AAF than with the secondary region A2 (AAF: 15%, A2: 10%). Finally, functionally-identified auditory areas were overall smaller than the atlas areas, and most of the atlas areas fell outside of the functionally-identified auditory areas of individual animals (Figure 2F, gray shading; AuD: 73%, Au1: 50%, AuV: 68%). As a result, if atlas coordinates are used to guide experimental manipulations (such as viral injections or electrophysiological recordings), it is highly likely that the manipulation will miss the target functional areas (Figure 2F, targeting accuracy: Au1→A1: 28 ± 2%, Au1→primary regions: 44 ± 2%, AuV→A2: 10 ± 1%). These results held true even when we lowered the threshold for area boundary detection to identify larger functional areas (40% peak response amplitudes; Supplementary Figure 3). These data indicate that using population-based resources, such as the brain atlas, is inadequate to accurately target functional cortical areas, whose locations are highly variable across individuals.

**Figure 2.**
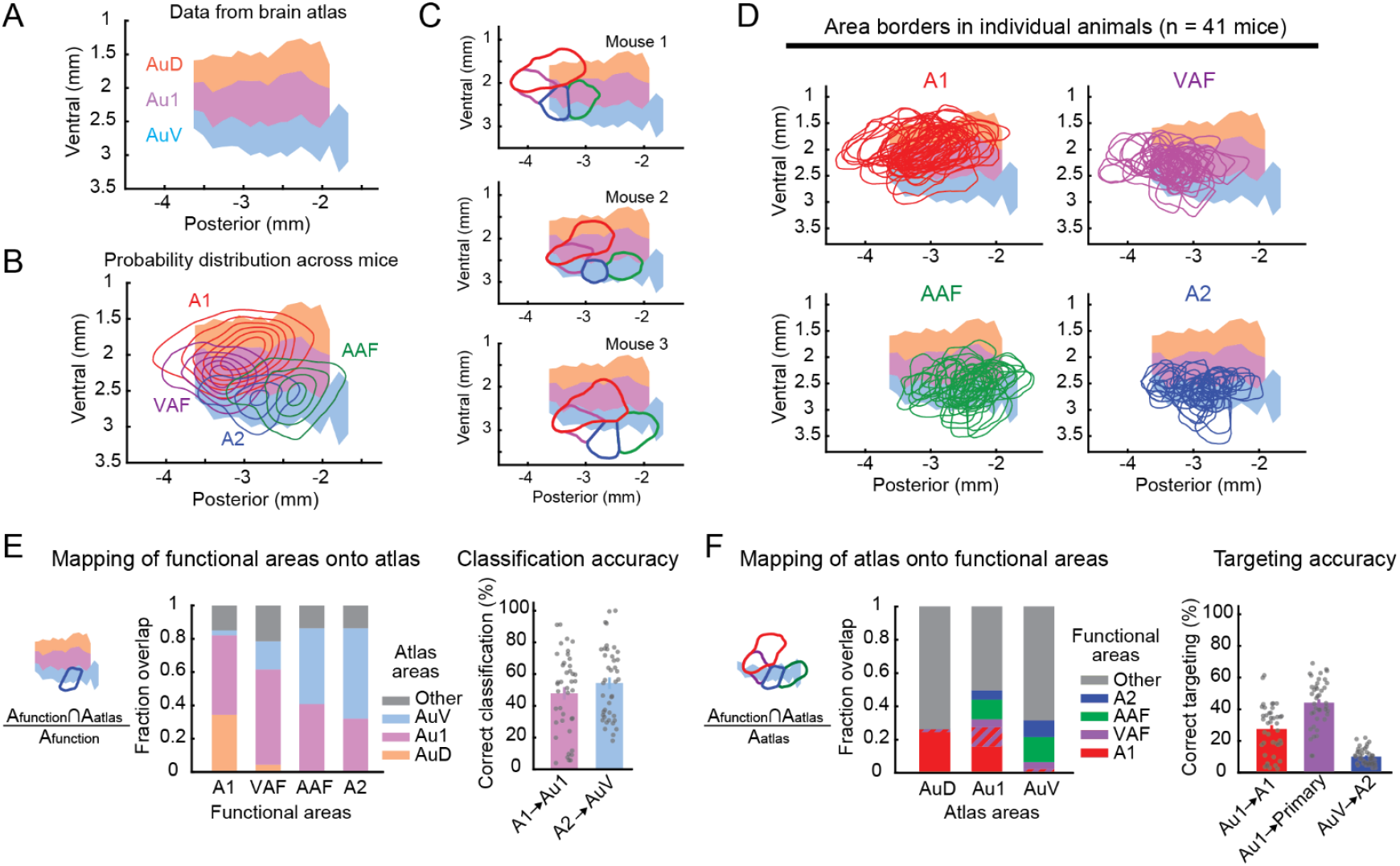
Stereotaxic coordinates based on the brain atlas cause substantial targeting errors. **(A)** A topographical surface map of auditory cortical areas based on the Paxinos Atlas. Coordinates are measured from bregma. **(B)** Probability distribution of functional auditory areas superimposed on atlas areas Au1, AuD, and AuV. Contours are 10% steps, starting at 10% (n = 41 mice). **(C)** Functionally-identified cortical area borders superimposed on the atlas map, showing inter-animal variability in their relationship to the atlas areas. The same three mice as Figure 1D. **(D)** Functionally-identified cortical area borders from all mice superimposed on the atlas map, shown separately for A1, VAF, AAF, and A2. **(E)** Left, fraction spatial overlap of functionally-identified areas with atlas areas. Right, classification accuracy showing the fraction of A1 contained within Au1 (A1→Au1) and A2 within AuV (A2→AuV). Each functional area tends to overlap with multiple atlas areas rather than being contained within a single area, resulting in only 48 ± 4% and 54 ± 4% accuracy (n = 41 mice; mean ± SEM). **(F)** Left, fraction spatial overlap of atlas areas with functionally-identified areas. Right, targeting accuracy showing the fraction of Au1 contained within A1 (Au1 → A1), Au1 within three primary areas (Au1 → Primary), and AuV within A2 (AuV → A2). Using stereotaxic coordinates to target functionally-defined auditory cortex results in only 28 ± 2%, 44 ± 2%, and 10 ± 1% accuracy, respectively (n = 41 mice; mean ± SEM).

### Stereotaxic targeting accuracy is low regardless of strain and sex

What biological factors contribute to the inter-animal variability in the absolute location of functional auditory cortices and the low accuracy of stereotaxic targeting? To investigate this, we first examined how functional maps generated from different mouse strains map onto the atlas-defined auditory cortex. We focused on the two most commonly used strains in auditory research: B6 and CBA. B6 is a source for many transgenic mouse lines and is used to generate brain atlases (Oh et al., 2014; Paxinos and Franklin, 2019, 2012; Q. Wang et al., 2020), and CBA is commonly used for auditory experiments due to its robustness against age-related hearing loss (Parham and Willott, 1988; Zheng et al., 1999). Using the coordinate system normalized to bregma–lambda distance, we found that functionally-identified auditory cortex centroids (mean of A1 3 kHz, A1 30 kHz, AAF 3 kHz, and A2 3 kHz centroids) in CBA mice were significantly more posterior than B6 mice (Figure 3A, B; B6: n = 14, CBA: n = 13 mice; AP, B6: −2.847 ± 0.248 mm, CBA: −3.132 ± 0.286 mm, p = 0.0094; DV, B6: 2.296 ± 0.145 mm, CBA: 2.311 ± 0.129 mm, p = 0.8651; mean ± SD, Wilcoxon rank sum test). The difference between strains was more prominent when we used the bregma-based absolute coordinates without scaling (Supplementary Figure 4), indicating the effectiveness of normalization for brain size. Nevertheless, even with size normalization and separation of strains, we still observed significant spatial variability in the area centroids, which often fell outside the atlas area boundaries. Stereotaxic targeting accuracy for primary and secondary areas were only 52 and 12%, respectively, in B6 mice, and the posterior shift of functional maps in CBA mice resulted in even lower accuracy (Figure 3C; B6 vs. CBA: p = 2.84×10^−6^, two-way ANOVA). We did not observe significant differences in area distributions between wild type B6 mice and transgenic strains, including PV-Cre×Ai9 and VGAT-Cre×Ai9 (Supplementary Figure 5).

**Figure 3:**
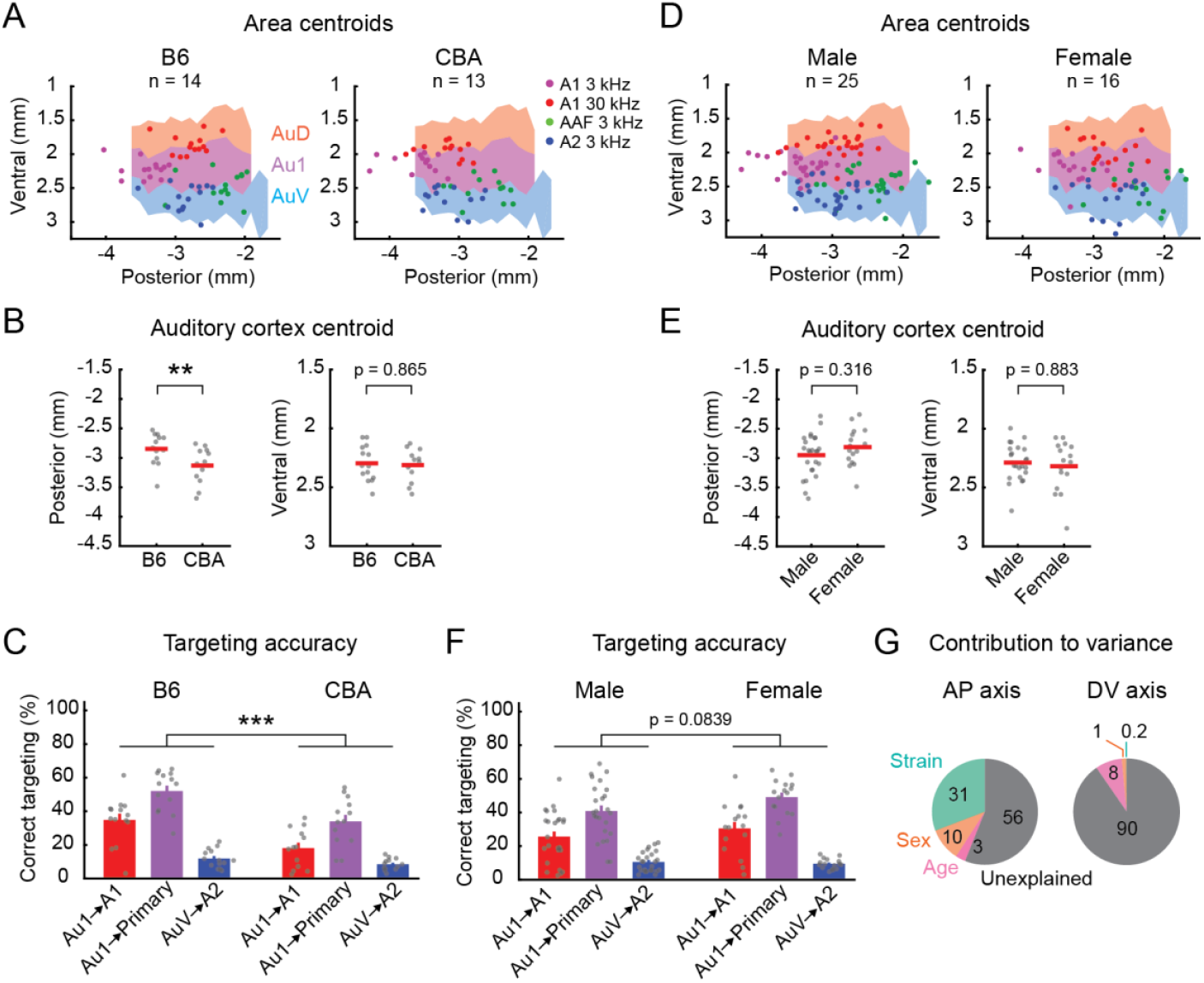
Strain and sex explain only a small fraction of spatial variability. **(A)** Distribution of functionally-identified frequency domain centroids shown separately for B6 (left, n = 14 mice) and CBA (right, n = 13 mice) mice. Scatter plots are superimposed on the atlas maps. **(B)** Scatter plots showing the posterior (left) and ventral (right) coordinates of functionally-identified auditory cortex centroids in individual B6 and CBA mice. Red lines are mean. *p = 0.0094, Wilcoxon rank sum test. **(C)** Accuracy of using atlas-defined stereotaxic coordinates to target functionally-identified areas in B6 and CBA mice. ***p = 2.84×10^−6^, two-way ANOVA. **(D**–**F)** Same as in (A–C) but for the comparison between males (left, n = 25 mice) and females (right, n = 16 mice). This dataset includes B6, CBA, PV-Cre×Ai9, and VGAT-Cre×Ai9 strains. **(G)** Pie charts displaying the percent contribution of strain, sex, and age to the total observed variance in the stereotaxic location of functionally-identified auditory cortex centroids. Data are shown separately for the anteroposterior (left) and dorsoventral (right) axes. The majority of the observed variance is unexplained by any of these three factors.

Next, we compared how functional maps generated from males and females mapped onto the atlas- defined auditory cortex. Although functionally-identified auditory cortex centroids in females had a slightly more anterior distribution compared to males, we observed no significant difference (Figure 3D, E; male: n = 25, female: n = 16 mice; AP, male: −2.948 ± 0.346 mm, female: −2.812 ± 0.308 mm, p = 0.3162; DV, male: 2.288 ± 0.147 mm, female: 2.319 ± 0.206 mm, p = 0.8831, mean ± SD; Wilcoxon rank sum test). Both sexes had significant variability across mice in how the functional areas mapped onto the atlas, which resulted in low, but similar, stereotaxic targeting accuracy (Figure 3F, male vs. female: p = 0.0839, two-way ANOVA). We did not observe a significant difference in stereotaxic locations between young (6–8 weeks old) and old (9–12 weeks old) mice (Supplementary Figure 6). To quantify the contribution of individual factors to the observed variability in stereotaxic locations, we performed an analysis of variance using strain, sex, and age as independent variables. While mouse strain contributed to a larger fraction of variance along the AP axis compared to sex and age (Figure 3G, AP, strain: 30.8%, sex: 10.1%, age: 3.0%; DV, strain: 0.2%, sex: 1.4%, age: 8.1%), most of the variance was unexplained by these factors (AP axis: 56.1%, DV: 90.4%), further confirming the seemingly random inter-individual variability in the cortical area locations. Taken together, these data show the necessity for functional mapping of cortical areas even in experiments using mice with uniform strain and sex. Furthermore, although B6 mice already display large variability in the location of functional cortical areas, using other strains needs further caution as it results in even larger deviation from the brain atlas.

### Au1 in the brain atlas overlaps with tone low-responsive auditory cortical areas

In addition to the tonotopic auditory areas we identified with intrinsic signal imaging, previous mapping studies identified non-tonotopic areas designated as dorsoposterior field (DP) as well as a center region (CTR) between A1 and AAF (Issa et al., 2017, 2014; Liu et al., 2019). In order to estimate the boundaries of these tone low-responsive areas, we used a lower threshold for automated area detection (20% peak response amplitudes). We defined DP as the tone low-responsive area dorsoposterior to A1 and CTR as the area surrounded by the tonotopic areas (Figure 4A). Interestingly, the distance between A1 and AAF was extremely variable, and therefore CTR was clearly present in some animals (see Figure 1C and Figure 4A) but was nearly absent in others (see Figure 1D). When we plotted the probability distribution of DP and CTR area masks across mice onto the Paxinos Atlas topographical map, these areas spanned across large areas of atlas-defined auditory cortex (Figure 4B). More specifically, DP fell partially onto AuD but was largely outside the atlas-defined auditory cortex, and CTR primarily mapped onto AuD and Au1 (Figure 4C). Therefore, tone low-responsive marginal auditory areas extend beyond the boundaries of the auditory areas defined in the brain atlas. When we quantified how the atlas areas map onto the functional areas, we found that AuD consists of a larger fraction of tone low-responsive areas than tonotopic areas (Figure 4D). Importantly, even though Au1 is commonly considered a primary cortex, it included a substantial fraction of CTR and non-auditory areas, which emphasizes that using Au1 to target primary cortex has a considerable chance of hitting tone low-responsive or non-responsive areas. Since tonotopic and non-tonotopic areas in mice show complex, interleaved distributions, our results emphasize the importance of accurately targeting specific cortical areas with functional mapping.

**Figure 4.**
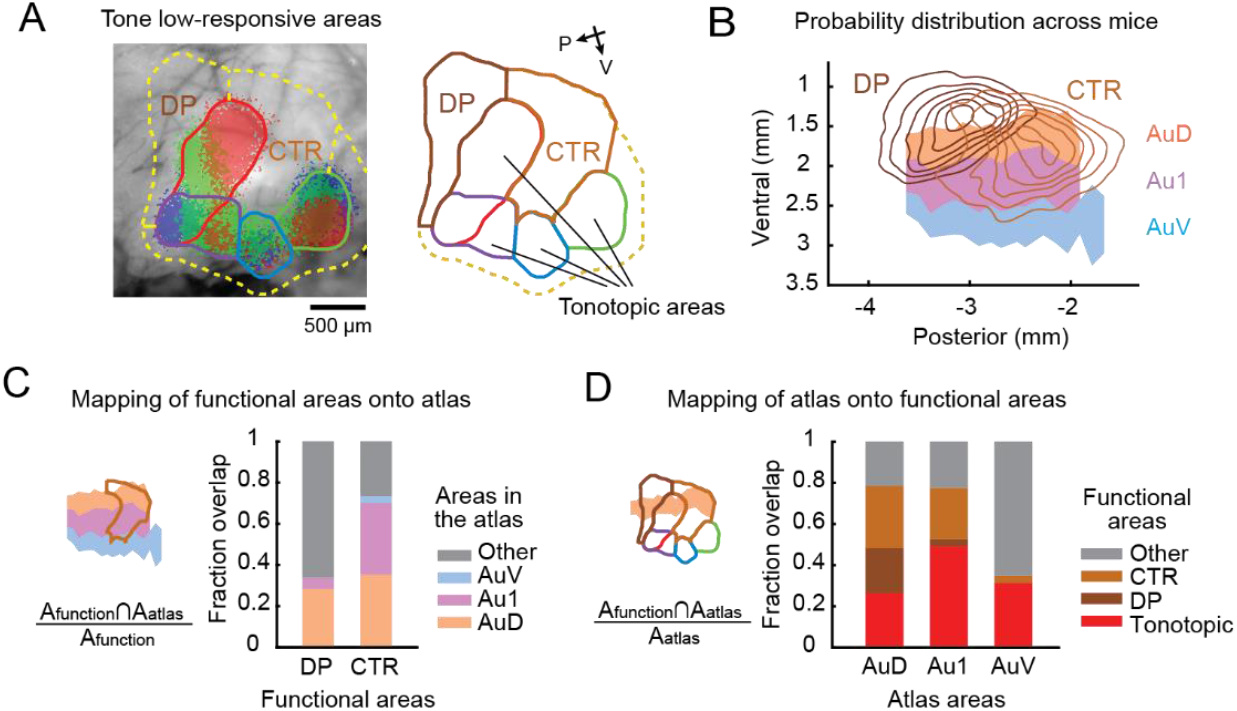
Au1 in the brain atlas contains tone low-responsive areas. **(A)** Automated segmentation of tone low-responsive dorsoposterior (DP) and center (CTR) areas using intrinsic signal imaging. Left, representative data showing the thresholded intrinsic signals and detected area borders superimposed on the cortical vasculature image. Yellow dotted line shows the boundary of tone low-responsive area. Right, extracted area borders, highlighting DP and CTR (see Methods for their definition). **(B)** Probability distribution of CTR and DP locations superimposed on the atlas map (n = 41 mice). Contours are 10% steps, starting at 10%. **(C)** Fraction spatial overlap of functionally-identified tone low-responsive areas with atlas areas. CTR has a substantial (35%) overlap with Au1. **(D)** Fraction spatial overlap of atlas areas with tone low-responsive areas. Au1 contains both tone low-responsive (29%) and non-auditory (22%) areas, highlighting the risk of using Au1 coordinates to target primary auditory cortex A1.

### Spatial variability is not simply due to suture irregularities but reflects inter-animal differences in cortical geography

We have found substantial inter-animal variability in the stereotaxic location of auditory cortex based on commonly used skull landmarks, bregma and lambda. Does this variability reflect irregular skull suture patterns across mice? Alternatively, do functional cortical area locations vary within the brain geography? To address this, we used an alternative approach to determine the stereotaxic coordinates of functionally-identified cortical areas without relying on skull landmarks. We took advantage of a previous histology dataset where we marked A2 with fluorophores by functionally-targeted craniotomies based on intrinsic signal maps (n = 20 mice). These data include both A2-targeted recordings, with the recording site marked by DiI or DiO, as well as A2-targeted viral injections. After slicing coronal brain sections, we identified the stereotaxic coordinates of the dye deposit by comparing the brain morphology with the Paxinos Brain Atlas (Figure 5A, B, Supplementary Figure 7). The knowledge of relative location between the craniotomy and the A2 border based on intrinsic signal imaging allowed us to determine A2 boundaries on the topographic atlas map. Consistent with our results using the stereotaxic coordinates based on skull landmarks (Figure 1 and Figure 2), the absolute location of A2 centroids showed large inter-animal variability on both the AP and DV axes (Figure 5C). The identified A2 borders overlapped the most with AuV (57%), while there was also substantial overlap with Au1 (28%) and temporal association area (TeA: 15%) (Figure 5C–E). Most strikingly, the accuracy of targeting A2 using AuV coordinates was only 9 ± 1% (Figure 5E), consistent with our results using skull landmarks (10 ± 1%; Figure 2F). These data showed a similar mismatch between the functional maps and the brain atlas regardless of whether we used skull landmarks or gross brain morphology to determine stereotaxic coordinates. Therefore, the spatial variability of cortical areas is not simply attributed to irregular suture patterns, but rather suggests that the relative positioning of functional cortical areas within the brain (cortical geography) varies across animals. Supporting this idea, we found high variability in the parameters describing the relative locations between auditory cortical areas in a large dataset of intrinsic signal imaging from our previous studies, which cannot be explained by global scaling or transposition (n = 300 mice; Figure 6A, B). Interestingly, the distance between A1 and AAF borders showed marked variability, explaining the inconsistency in identifying CTR across animals (Figure 6B). Therefore, cortical geography is highly variable across individuals at both the local (relative location between auditory areas) and global (positioning of the overall auditory cortex within the brain) scale. Taken together, our results demonstrate that functionally-defined cortical areas are not spatially fixed within the brain geography and vary in size, shape, and relative locations across animals. Functional mapping of individual mice is therefore required for accurately targeting specific areas for manipulation, as generalization across populations fails to capture inter-animal variability.

**Figure 5:**
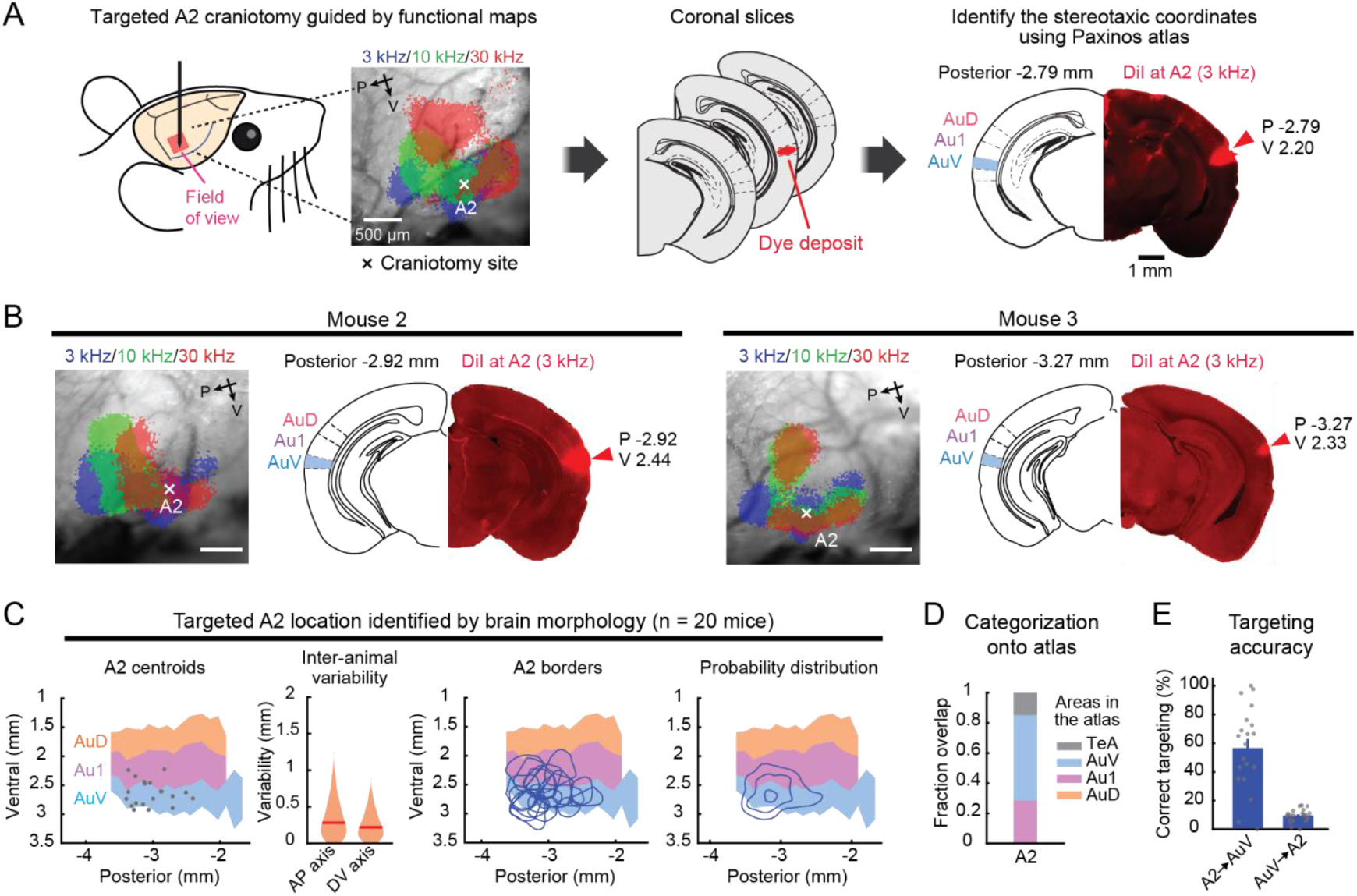
A2 location variability is not simply due to suture irregularities but reflects inter-animal differences in cortical geography. **(A)** Experiment and analysis setup. A2 craniotomy location was recorded during surgery. After coronal sectioning, the section with the strongest fluorescence was identified. Stereotaxic locations of the dye deposit and A2 area boundary were determined using the Paxinos Brain Atlas (details in Supplementary Figure 7). **(B)** Representative data for the determination of dye deposit locations in two additional mice. Left, Intrinsic signal imaging maps showing craniotomy sites. Right, coronal sections with the strongest fluorescence and their corresponding atlas schematics shown side-by-side. **(C)** Distributions of A2 area centroids, boundaries, and their probability distribution from all mice superimposed on the atlas map (n = 20 mice; breakdown of genotypes shown in Methods). Contours are 10% steps, starting at 10%. Violin plots show inter-animal variability of A2 centroid locations along anteroposterior (left) and dorsoventral (right) axes. n = 190 mouse pairs. Red lines are median. **(D)** Fraction spatial overlap between A2 as determined by dye deposit and atlas areas. **(E)** Left, classification accuracy showing the fraction of A2 contained within AuV (A2→AuV). Right, targeting accuracy showing the fraction of AuV contained within A2 (AuV→A2). These measurements without using suture landmarks are similar to those in Figure 2E and 2F, indicating that the variability is not explained by suture irregularity. Atlas section schematics were drawn based on the Paxinos and Franklin’s Mouse Brain Atlas (Paxinos and Franklin, 2019).

**Figure 6:**
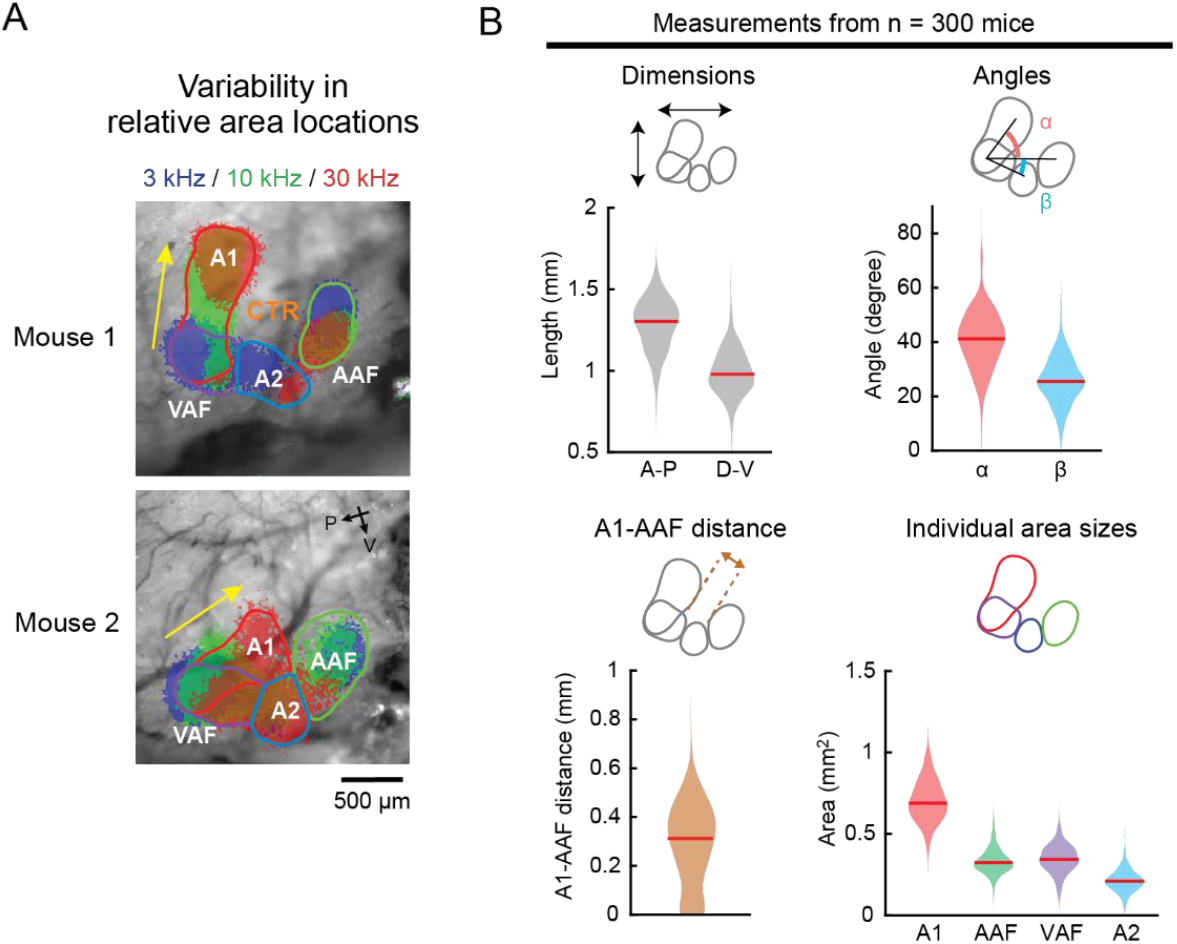
Variability in the local geography of functionally-identified auditory cortical areas. **(A)** Thresholded intrinsic signals and detected area borders of two representative mice showing variable relative positions of functional areas. Yellow arrows indicate the directions of A1 tonotopic axes. Mouse 1 shows a dorsoventrally oriented A1 axis with a clear CTR region, whereas Mouse 2 has an anteroposteriorly oriented A1 axis with little room for CTR. **(B)** Violin plots showing the variability in the local geography measurements of functional auditory cortical areas (n = 300 mice; breakdown of genotypes shown in Methods). Top left, variability in the size of the overall auditory cortex in anteroposterior and dorsoventral dimensions. Top right, variability in the relative positioning of functional areas measured by the angle between A1 axis and A1–AAF axis (α), and between A1–A2 axis and A1–AAF axis (β). Bottom left, variability in the distance between A1 and AAF borders, which indicates the dimension of a tone low-responsive CTR region. Bottom right, variability in the area of functionally-identified A1, AAF, VAF, and A2. Red lines are median.

## Discussion

### Inter-animal spatial variability of auditory cortical areas

In this study, we used intrinsic signal imaging to directly measure the stereotaxic locations of functionally-identified auditory cortices from a large group of animals. We found marked inter-animal variability, with the mean and maximum offsets as large as 410 and 1,584 μm for the AP axis, and 215 and 1,004 μm for the DV axis. As these values are comparable to the width of individual functional auditory areas, our results suggest a high probability of targeting errors when using stereotaxic coordinates. The observed inter-animal variability likely reflects a combination of multiple factors, such as human errors in stereotaxic measurements, absolute brain size differences across strains (Paxinos et al., 1985; Wahlsten et al., 1975), irregular suture patterns (Blasiak et al., 2010; Whishaw et al., 1977; Zhou et al., 2020), and variability in the relative positioning of functional areas within the cortex (cortical geography) (Garrett et al., 2014; Waters et al., 2019). We concluded that heterogeneity of cortical geography dominates the inter-animal variability in the stereotaxic coordinates for the following three reasons. First, we directly quantified human-related experimental variability by comparing stereotaxic markings across experimenters and found it to be only a minor factor (Figure 1F), suggesting biological rather than human origins of variability in our conditions. Nevertheless, we do not exclude the possibility that human errors further deteriorate stereotaxic targeting in other studies, where criteria for locating suture landmarks may not be standardized across researchers. Second, explicit attributes of animals that may affect brain sizes, such as strain, sex, and age, explained less than half of the total spatial variability (Figure 3). The negligible effect of age is consistent with the stable cranial size within the age range used in our study (6–12 weeks old) (Vora et al., 2016). We found significant inter-strain, but not inter-sex, differences in stereotaxic coordinates (Paxinos et al., 1985; Wahlsten et al., 1975). Importantly, although the selective inclusion of B6 mice and normalization of AP coordinates somewhat reduced inter-animal variability, the remaining variability was still evident and caused substantial errors in stereotaxic targeting. Therefore, suture landmarks alone are insufficient for targeting functional cortical areas, regardless of whether coordinates are scaled to brain sizes. Finally, in an independent set of experiments that determined A2 stereotaxic coordinates without using suture landmarks, we found spatial variability comparable to the bregma-based method (Figure 5), which directly demonstrates the heterogeneity of cortical geography across individuals. This conclusion was further supported by our large-scale (n = 300 mice) imaging dataset showing considerable variability in size, orientation, and relative locations of auditory cortical areas (Figure 6). Together, these results demonstrate that functionally-defined cortical areas are not spatially fixed within the brain geography, which could lead to substantial stereotaxic targeting errors unless functional mapping is conducted in individual animals.

This surprisingly large variability of cortical geography is consistent with previous reports which quantified the relative locations between visual cortical areas across mice (Garrett et al., 2014; Waters et al., 2019). Although these studies did not measure the absolute stereotaxic coordinates from bregma, they found up to 1 mm variability in relative cortical locations, in agreement with our observation in the auditory areas. The variability of cortical geography even within uniform genetic backgrounds raises an interesting possibility that developmental experience may influence the relative location of sensory cortices. Numerous previous studies have found plasticity in A1 spatial organization depending on sound experience during the critical periods of development (Barkat et al., 2011; De Villers-Sidani et al., 2007; Han et al., 2007; Insanally et al., 2009; Kim and Bao, 2009; Zhang et al., 2001). Although all our mice were raised in a uniform low-noise sound environment, experiences unique to individual animals, such as interactions with their littermates and parents, could have influenced the size and location of auditory cortical areas. Since experience-dependent cortical reorganization is also established in visual and somatosensory systems (Simons and Land, 1987; Wiesel and Hubel, 1963), global cortical geography may vary as a result of developmental experience across sensory modalities. In the future, it would be of interest to investigate how controlled manipulation of sensory environments alters the absolute stereotaxic locations of primary and secondary sensory cortical areas.

### Comparison between the atlas map and functional maps

In the commonly used mouse brain atlases, the overall location of auditory cortex was determined according to the immunostaining of neurofilament using SMI-32 antibody (Oh et al., 2014; Paxinos and Franklin, 2019, 2012; Wang et al., 2011). Within the auditory cortex, primary Au1 is surrounded by two higher-order areas, AuD and AuV, which run in parallel rostrocaudally. As described in the preface of the Paxinos Brain Atlas, these boundaries and nomenclature were adopted from the rat brain atlas (Paxinos and Franklin, 2012), and they likely had their origin in the “core and belt” structure in the auditory cortex of primates (Hackett et al., 1998; Merzenich and Brugge, 1973). Expression patterns of calcium-binding proteins, such as parvalbumin, calbindin, and calretinin, were used to help separate the primary and secondary cortices (Cruikshank et al., 2001; Jones et al., 1995). However, their gradational distributions pose challenges in drawing exact area boundaries, which likely resulted in the mismatches between the Paxinos and the Allen Institute brain atlases.

In contrast, functional mapping has revealed a more complex spatial organization of the mouse auditory cortex, using both electrophysiological (Guo et al., 2012; Joachimsthaler et al., 2014; Stiebler et al., 1997) and optical methods (Aponte et al., 2021; Issa et al., 2014; Kline et al., 2021; Liu et al., 2019; Romero et al., 2020; Tsukano et al., 2015). There is a general agreement that there are four tonotopic subregions, A1 (including the ultrasound field), VAF (or the ventral branch of A1), AAF, and A2, each of which has its unique direction of tonotopic gradient. In addition to these tonotopic regions, the existence of at least two non-tonotopic, tone low-responsive areas, DP and CTR, have been proposed. Although there is still ongoing argument regarding the definition of individual area borders and the existence of CTR (Romero et al., 2020), an important consensus is that the primary auditory cortices extend dorsoventrally and are therefore unlikely to be contained within the atlas Au1. These observations suggest that the simple core–belt structure in the brain atlases does not represent the actual complexity of the functional map.

Our direct measurement of the stereotaxic coordinates of functionally-identified cortical areas provided critical insights into the relationship between the atlas map and functional maps. We found that the brain atlas accurately described the population-averaged location of functionally-identified auditory cortex as a whole (Figure 2B). However, when we considered individual mice, the functional auditory cortex was more compact than the atlas auditory cortex, and the spatial relationship between the atlas and functional areas was highly variable (Figure 2C, D). An example of a small and ventrally shifted auditory cortex can also be found in a previous study that marked the edges of the functionally-identified auditory cortex (Romero et al., 2020). Since the Paxinos Brain Atlas was constructed using brains from 26 mice (Paxinos and Franklin, 2019), it is reasonable that the inter-animal variability in these brains was averaged out to generate a broader population-level distribution. Consequently, each subdivision in the atlas auditory cortex pools multiple functional areas, including not only primary and secondary auditory cortices but also surrounding non-auditory regions (Figure 2F). Therefore, using stereotaxic coordinates based on the brain atlas obscures experimental data by erroneously merging results from multiple functional areas, emphasizing the importance of functional mapping in individual animals.

Au1 in the brain atlas has been commonly used as a proxy for the primary auditory cortex. Our data showed that functionally-identified A1 occupies only 28% of Au1, and even if we combined three primary areas, A1, VAF, and AAF, they covered only 44% of Au1. The rest of Au1 was largely divided into tone low-responsive CTR (25%) and non-auditory areas (22%). Whether CTR is a part of the primary area AAF or a higher-order cortex has been debated. Traditionally, the tonotopic axis of AAF was drawn from the low-frequency pole of AAF toward the high-frequency pole of A1 (Guo et al., 2012; Romero et al., 2020; Stiebler et al., 1997). However, a tone low-responsive domain between A1 and AAF (CTR) has been observed repeatedly in mice (Aponte et al., 2021; Ceballo et al., 2019; Honma et al., 2013; Issa et al., 2017, 2014; Kline et al., 2021; Liu et al., 2019; Tsukano et al., 2015) and rats (Polley et al., 2007; Profant et al., 2013). This CTR area was reported to respond preferentially to complex sounds over pure tones (Honma et al., 2013; Issa et al., 2017, 2014; Tsukano et al., 2015). Furthermore, faint SMI-32 staining and a high fraction of inputs from the secondary thalamus (Honma et al., 2013; Tsukano et al., 2015) suggested that CTR may be one of the higher-order auditory cortices. In the current study, we found a surprising heterogeneity in the distance between functionally-defined A1 and AAF borders (Figure 6), suggesting that at least a part of the previous controversy may be attributed to inter-animal variability. For example, CTR is likely positioned dorsal to AAF in animals with adjacent A1–AAF (Mouse 3 in Figure 1D), while it may intrude into the area between A1 and AAF in mice with a dorsoventrally-oriented A1 (Figure 1C). Future studies will elucidate more detailed characteristics of CTR and factors influencing its size and shape. Regardless of whether CTR is primary or secondary, a critical conclusion here is that the atlas Au1 includes a substantial fraction of tone low-responsive and non-auditory areas.

The most striking dissociation between our functional mapping and the common usage of the brain atlases was found in the relationship between AuV and A2. In contrast to the general belief that AuV represents a secondary A2 area (although it is controversial whether A2 is truly a secondary cortex (Ohga et al., 2018; Romero et al., 2020)), functionally-identified A2 occupied only 10% of the atlas AuV (Figure 2F). This mismatch was mainly due to A2’s compact size and limited extension along the AP axis (Issa et al., 2014; Kline et al., 2021; Liu et al., 2019; Ohga et al., 2018; Romero et al., 2020). This conclusion was robust even when we expanded the A2 boundary by lowering the area detection threshold (Supplementary Figure 3). Indeed, AuV had more overlap with the primary AAF (15%), and the combined primary areas (A1, VAF, and AAF) occupied twice the area of AuV (22%) than A2. Moreover, 68% of AuV fell outside the functionally-identified auditory cortical areas, suggesting its contamination from the adjacent non-auditory areas, such as the temporal association area and secondary somatosensory area. Together with the fact that functional A1 occupies 27% of AuD, these results indicate that the general categorization of Au1 as a primary area and AuD and AuV as secondary areas is highly prone to errors.

### Functional mapping for the investigation of cortical area-specific roles

We found surprisingly low accuracy of targeting functional cortical areas based on stereotaxic coordinates. The estimated error fraction reached as high as 56% in using Au1 to target primary auditory areas and 90% in using AuV to target A2. These errors were due not only to the simple area segmentation scheme in the brain atlas but also to the marked inter-animal variability in cortical geography. Although this spatial variability may not pose a substantial problem in large areas such as primary visual cortex, it makes stereotaxic targeting extremely challenging in small areas, such as the auditory cortex and secondary visual cortex. For example, neurons preferring temporally coincident multi-frequency sounds are spatially clustered in a highly restricted area within A2 (Kline et al., 2021), and their identification and selective manipulation were possible only with functional mapping. Recent efforts in using functional mapping to focally manipulate specific areas have proven their power in dissecting the area-specific roles in the auditory cortex (Ceballo et al., 2019; Kline et al., 2021). Since there are no acceptable cytoarchitectural landmarks that allow post hoc histological identification of functional areas in the neocortex, performing functional mapping in each animal is the only way to target cortical areas without ambiguity. The same rule applies to the interpretation of histology data, such as annotating tracing and gene expression data to specific functional areas. We recommend that researchers perform functional mapping in each mouse and mark the cortical area locations before dissecting the brains for histology, similar to the approach in Figure 5 and recent studies (Romero et al., 2020; Tsukano et al., 2016). The area selectivity achieved with this extra step (which takes no more than 1.5 hours, including the entire surgery) helps researchers obtain reproducible data without erroneously pooling multiple functional areas, thus saving time and reducing the number of animals used in the end.

Which methods are the most suitable for functional mapping of cortical areas? The ideal method should be quick, easy, non-invasive, and broadly applicable to various experiments. Optical imaging is thus a better choice than electrophysiological mapping, which typically requires multiple penetrations with a large craniotomy. In particular, intrinsic signal imaging (Aponte et al., 2021; Bathellier et al., 2012; Ceballo et al., 2019; Grinvald et al., 1986; Kalatsky et al., 2005; Kato et al., 2017, 2015; Kline et al., 2021; Nelken et al., 2004) is an ideal candidate for the following four reasons. First, transcranial intrinsic signal imaging keeps the skull and cortex intact as it does not require invasive procedures such as craniotomy, skull thinning, dye infusion, or gene transfection. Second, the intrinsic signal is robust enough that a few trials are typically enough to visualize coarse maps, thus enabling quick mapping that can be easily combined with other surgical procedures. Third, intrinsic signal imaging does not require extrinsic genes, reducing the labor of breeding transgenic mice. Lastly, intrinsic signal imaging is robust against background fluorescence in the cortex as it uses light reflection and not fluorescence. It is highly compatible with modern neuroscience techniques, which often take advantage of genetically encoded fluorescent proteins. Other imaging techniques, such as macroscopic GCaMP calcium imaging (Issa et al., 2017, 2014; Liu et al., 2019; Romero et al., 2020) and autofluorescence imaging (Shibuki et al., 2003; Tsukano et al., 2015), may also be useful when extrinsic fluorescent gene expression is unnecessary. Nevertheless, we generally recommend intrinsic signal imaging, as it offers the largest flexibility with experiment designs. In this manuscript, we provide a detailed protocol for intrinsic signal imaging to facilitate the adoption of functional mapping by cortical researchers (Supplementary Protocol). We hope that this study helps the field advance our understanding of information processing in hierarchically organized cortical streams by providing a more reliable reference frame to dissect area-specific functions.

## Materials and Methods

### Animals

Mice were 6–12 weeks old at the time of experiments. Mice were acquired from Jackson Laboratories: C57BL/6J; CBA/J; Slc32a1^tm2(cre)Lowl^/J (VGAT-Cre); Pvalb^tm1(cre)Arbr^/J (PV-Cre); Gt(ROSA)26Sor^tm9(CAG-tdTomato)Hze^/J (Ai9). For Figure 6, data from our previous studies were reanalyzed (Aponte et al., 2021; Kline et al., 2021; Onodera and Kato, 2022). For this dataset, additional mice were acquired from Jackson Laboratories (Sst^tm2.1(cre)Zjh^/J (Sst-Cre)) and MMRRC (Tg(Rbp4-Cre)KL100Gsat/Mmucd (Rbp4-Cre); Tg(Tlx3-Cre)PL56Gsat/Mmucd (Tlx3-Cre)). Both female and male animals were used and housed at 21°C and 40% humidity with a reverse light cycle (12–12h). All experiments were performed during their dark cycle. All procedures were approved and conducted in accordance with the Institutional Animal Care and Use Committee at the University of North Carolina at Chapel Hill as well as the guidelines of the National Institutes of Health.

### Sound Stimulus

Auditory stimuli were calculated in Matlab (Mathworks) at a sampling rate of 192 kHz and delivered via a free-field electrostatic speaker (ES1; Tucker-Davis Technologies). Speakers were calibrated over a range of 2–64 kHz to give a flat response (±1 dB). For functional mapping with intrinsic signal imaging, 3, 10, and 30 kHz pure tones (75 dB SPL, 1-s duration) were presented at a 30-s interval. Pure tone stimuli had a 5-ms linear rise-fall at their onsets and offsets. Stimuli were delivered to the ear contralateral to the imaging site. Auditory stimulus delivery was controlled by Bpod (Sanworks) running on Matlab.

### Marking of stereotaxic reference points

Prior to intrinsic signal imaging, three stereotaxic reference points were marked on the skull with black ink to allow for the integration of functionally-identified auditory cortices into the stereotaxic coordinate system. Mice were anesthetized with isoflurane (0.8–2%) vaporized in oxygen (1 L/min), kept on a feedback-controlled heating pad at 34–36 °C, and placed on a stereotaxic frame (Kopf Instruments Model 1900). The mouse was secured with ear bars and a palate bar, and the scalp and the muscle overlying the right auditory cortex were either removed or pushed aside. A sharpened metal needle attached to a three-axis motorized manipulator (Scientifica IVM) was used to level bregma and lambda by rotating the head around three axes (roll, yaw, and pitch). We followed the definitions of bregma and lambda in the Paxinos Brain Atlas (Paxinos and Franklin, 2019, 2012), except that steep curves in the coronal suture near the midline were ignored by drawing fit lines. Stereotaxic marking was made for the following three coordinates relative to bregma: (Posterior, Ventral) = (−2.5, 1.5), (−3.5, 1.5), and (−3.5, 2.0) (in millimeters). In a small subset of mice, coordinates of (−2.5, 1.0), (−3.5, 1.0), (−3.5, 2.0) (n = 5 mice) or (−2.5, 1.2), (−3.5, 1.2), (−3.5, 2.0) (n = 2 mice) were used. The distance between bregma and lambda (BLdist) was measured in each mouse, and the coordinates of the reference points were scaled by a factor of BLdist/4.2 (4.2 mm refers to the standard BLdist for adult B6 males in the Paxinos Brain Atlas) to account for skull size differences. The tip of a metal needle was painted with black ink, and dots were made at three reference points by gently touching the skull surface with the needle. Following ink marking, intrinsic signal imaging was performed (see below). The ink markings were visualized in a brain surface image captured in the same field of view as the intrinsic signal, which allowed for direct comparison between the functional map and stereotaxic locations. In a subset of mice, to quantify human-related errors, the mouse was removed from the stereotaxic frame after imaging, and a second experimenter realigned the head and marked the stereotaxic reference points with ink. For each of the three reference points, the distance between dots made by two experimenters was calculated along the AP and DV axes. The distances were averaged across three reference points to give one data point of the human error value per animal. Various combinations of three experimenters performed ink markings.

### Intrinsic signal imaging (see Supplementary Protocol for more detail)

Intrinsic signal images were acquired using a custom tandem lens macroscope (composed of Nikkor 35 mm 1:1.4 and 135 mm 1:2.8 lenses) and a 12-bit CMOS camera (DS-1A-01M30, Dalsa) placed in a sound isolation chamber (Gretch-Ken Industries). After marking stereotaxic reference points, a custom-designed stainless steel head bar was attached to the skull using a small amount of dental cement. Mice were injected subcutaneously with chlorprothixene (1.5 mg/kg body weight) prior to imaging and kept under isoflurane anesthesia (0.8%). The brain surface was imaged through the skull kept transparent by saturation with phosphate-buffered saline. Images of surface vasculature were acquired using green LED illumination (530 nm), and intrinsic signals were recorded (16 Hz) using red illumination (625 nm) with a custom Matlab program. Images were acquired at 717 × 717 pixels (covering 2.3 × 2.3 mm^2^) or 1024 × 1024 pixels (covering 3.3 × 3.3 mm^2^). Each trial consisted of a 1-s baseline followed by a 1-s pure tone stimulus (75 dB SPL; 3, 10, or 30 kHz) and a 30-s intertrial interval. Images during the response period (0.5–2 s from the sound onset) were averaged and divided by the average image during the baseline. Images were Gaussian filtered (σ = 2 pixels) and averaged across 10–20 trials for each sound using IO and VSD Signal Processor Plugin on Fiji software (https://imagej.net/software/fiji/; https://murphylab.med.ubc.ca/io-and-vsd-signal-processor/) (Harrison et al., 2009). The resulting images were deblurred with a 2-D Gaussian window (σ = 200 μm, which corresponds to 68 pixels) using the Lucy-Richardson deconvolution method (Issa et al., 2014; Romero et al., 2020) to generate a trial-averaged response intensity map. Individual auditory areas including A1, AAF, VAF, and A2 were identified based on their characteristic tonotopic organization. For visualization of the functional maps in the figures, signals were thresholded independently for each sound (see the left panel of Figure 1C).

### Semiautomated sorting of area boundaries

For each sound frequency, frequency domains of four tonotopic areas (A1, AAF, VAF, and A2; see the middle panel of Figure 1C) were semiautomatically determined, using the trial-averaged response intensity map in the following steps. First, images were visualized with high thresholds to facilitate segregation and identification of the response centers for four areas. Guided by this thresholded map, coarse locations of frequency domains for four areas (seed ROIs) were manually drawn. The centroid of each frequency domain was determined as the mean of the two locations: peak response amplitude point and the center of mass point within the seed ROI. Next, around this domain centroid, the frequency domain mask was determined as the pixels whose signal intensity exceeded a fixed threshold of 60% of the peak amplitude within the seed ROI. When the masks of the same frequency domains overlapped between two cortical areas, a dividing line was drawn such that the distances from the dividing line to the two domain centroids were proportional to the peak response amplitudes in individual domains. As an exception, we allowed overlap between the 3 kHz domains of A1 and VAF since we observed convergence of these two areas at their low-frequency poles in most animals (Aponte et al., 2021; Issa et al., 2014; Kline et al., 2021; Liu et al., 2019; Romero et al., 2020). For Supplementary Figure 3, a lower threshold of 40% was used. Note that the area borders drawn in this method are largely independent of the initial selection of the seed ROIs, as long as the seed ROIs include the peak response amplitude points and are segregated from each other enough. Once the frequency domain masks were determined for three frequencies, these masks were combined to create the area masks for A1, AAF, VAF, and A2. After joining the binarized domain masks for three frequencies, the boundaries were smoothened with opening and closing operations (using disk-shaped elements of 30 and 150 pixels radius, respectively). In some mice, A2 and VAF lacked clear responses to one or two of the three frequencies tested, and therefore only the frequency domains for the responsive sounds were used. Finally, if the masks for two cortical areas overlapped with each other, a dividing line that passed the two intersection points was drawn. This additional step of overlap removal was necessary since the initial determination of frequency domain masks did not restrain the overlap between domains for different frequencies. Again, we allowed overlap between A1 and VAF masks. For identification of DP and CTR, individual frequency domains were determined in the same manner as described above, except that a 20% threshold was used to identify tone low-responsive pixels. After frequency domains from all sounds and cortical areas were combined together, the boundary of the combined mask was smoothened with opening and closing operations (using disk-shaped elements of 30 and 150 pixels radius, respectively). The exclusion of four tonotopic area masks from this combined mask left a single mask of tone low-responsive marginal areas. Within this tone low-responsive area, DP was defined as the area dorsal to the most posterior point of A1 and posterior to the most dorsal point of A1. Also within the tone-low-responsive area, CTR was defined as the area surrounded by 1) borders of A1, VAF, A2, and AAF, 2) a DV line that passes the most dorsal point of A1, and 3) a fit line to the AAF tonotopic axis.

### Integration of area masks into the stereotaxic coordinate systems

The auditory area centroids and masks were integrated into the stereotaxic coordinate system, using three stereotaxic reference points marked on the skull. Locations of the ink markings were manually identified in a magnified skull surface image, which was taken at the same location as the intrinsic signal imaging field of view, using Illustrator software (Adobe Inc). The three reference points at (P, V) = (−2.5, 1.5), (−3.5, 1.5), (−3.5, 2.0) (hereafter, referred to as marks 1, 2, and 3) allowed the identification of the AP and DV axes of stereotaxic coordinates. In some animals in which the AP (marks 1→2) and DV (marks 2→3) lines did not cross perpendicularly due to the slight skull curvature, a foot of the perpendicular line from mark 1 onto mark 2–3 line was used as the adjusted mark 2. The auditory area centroids and masks were rotated to make the AP and DV axes parallel to the x and y axes of the plot, respectively. As the markings were made using the coordinates scaled by a factor of BLdist/4.2 (see Marking of stereotaxic reference points section), the distances between marks 1–2 and 2–3 pairs were 1 mm × BLdist/4.2 and 0.5 mm × BLdist/4.2, respectively. Using these distances, scaling factors for converting intrinsic signal image pixels to micrometers were calculated separately for AP and DV axes. After the rotation and scaling, the absolute stereotaxic coordinates of the area centroids and masks from bregma were determined.

Three distinct coordinate systems were used throughout the study to describe the stereotaxic locations:

1. Absolute coordinates from bregma: using the raw absolute coordinates calculated above, therefore (P_bregma_, V_bregma_) = (0, 0) and (P_lambda_, V_lambda_) = (−BLdist, 0);
2. Absolute coordinates from lambda: using the raw absolute coordinates, but shifting the space such that (A_lambda_, V_lambda_) = (0, 0) and (A_bregma_, V_bregma_) = (BLdist, 0);
3. B–L Normalization: scaling the AP coordinates by a factor of BLdist/4.2 such that (P_bregma_,V_bregma_) = (0, 0) and (P_lambda_, V_lambda_) = (−4.2, 0).

P, A, and V refer to the posterior, anterior, and ventral coordinate values. The B–L normalization system was used throughout the study except for Supplementary Figures 1, 4, and 5, since this method minimized the inter-animal variability in auditory area locations. DV coordinates were not scaled, as the scaling rather increased the inter-animal variability (Supplementary Figure 1B).

### Generation of a topographical map of Paxinos Brain Atlas auditory areas

The topographical cortical surface map of the atlas auditory areas was generated using the Paxinos and Franklin’s Mouse Brain Atlas, fifth edition (Paxinos and Franklin, 2019). The DV coordinates of the dorsal and ventral edges of Au1, AuV, and AuD were extracted from 17 atlas brain sections between −1.67 to −3.63 mm posterior from bregma (Supplementary Figure 2, left and middle panels). There was a discontinuity in the AuV map at P = −1.91 mm, which was likely a mistake in the brain atlas. Therefore, at this location, ‘Au1’ in the brain atlas was divided equally into Au1 and AuV to achieve a smooth transition. After obtaining the dorsal and ventral edges of individual areas in all brain sections, these edges were connected across 17 sections along the AP axis (Supplementary Figure 2, right).

### Quantification of inter-animal variability

Data analyses of inter-animal spatial variability were conducted using the functional area centroids and masks integrated into the stereotaxic coordinates as described above. The probability distribution of functional auditory areas was visualized by overlaying area masks from all mice, applying a circular averaging filter (100 μm radius), and displaying the contour at 10% increment, starting from 10% (Figures 2B, 4B, and 5C). The mapping of functional areas onto the atlas was determined by calculating the intersection between each functional area and each atlas area, divided by the area of the functional region. The averaged data across all mice were displayed as bar graphs (Figure 2E, 4C, and 5D). Classification accuracy was calculated as the fraction of A1 contained within Au1 (A1→Au1) and A2 contained within AuV (A2→AuV) for individual animals. These values indicate the likelihood of an event within a functional area being correctly classified into its corresponding atlas area. The mapping of atlas areas onto the functional areas was determined by calculating the intersection between each atlas area and each functional area divided by the area of the atlas region. The averaged data across all mice were displayed as bar graphs (Figure 2F and 4D). Targeting accuracy was calculated as the fraction of Au1 contained within A1 (Au1→A1), Au1 contained within any of the primary areas (Au1→Primary), and AuV contained within A2 (AuV→A2) for individual animals. These values indicate the likelihood of hitting the desired functional region with experimental manipulation if stereotaxic coordinates from the brain atlas are used for targeting. For the inter-strain, inter-sex, and inter-age comparisons, the auditory cortex centroid was determined as the center of four area centroids: A1 3 kHz, A1 30 kHz, AAF 3 kHz, and A2 3 kHz for each mouse. The contribution of biological factors to the total spatial variance was calculated using Matlab’s anovan function. Strain, sex, and age were used as variables, with only age as a continuous variable.

### Identification of stereotaxic coordinates for dye deposits in A2

The dye deposit dataset in A2 (Figure 5) was from our A2-targeting experiments, including mice in our published study (Kline et al., 2021). The fluorescence signals were either from 1) virus injections with the localized expression of fluorescent markers (AAV9.hsyn.Flex.ChrimsonR.tdTomato or AAV5.syn.EYFP; UNC Vector Core), 2) DiI/DiO coating on a silicone probe used for *in vivo* unit electrophysiology, or 3) injections of fluorophore-conjugated cholera toxin subunit B (CTB 488 or CTB 555; Invitrogen). These manipulations were all targeted to A2, guided by intrinsic signal imaging. After sectioning the brain at 40 μm thickness, the section with the strongest fluorescence was identified as the center of the marking and used to identify stereotaxic coordinates with the Paxinos Brain Atlas (Supplementary Figure 7). The posterior coordinate of the dye deposit was determined by identifying the atlas section with the corresponding morphology. Since the cortex in the histology sections was often vertically extended compared to the brain atlas, the ventral coordinate was determined by vertically scaling the histology section to the height of the atlas section (schematics in Supplementary Figure 7). This was achieved by calculating V = v_top_ + (v_bottom_ – v_top_) × b/a, where v_top_ and v_bottom_ represent the ventral coordinates of the top and bottom of the atlas section, and a and b represent the histology section height and the distance from the histology section top to the dye deposit center, respectively. The injection/probe insertion location within the intrinsic signal imaging map was recorded during the experiment. The relative location between the craniotomy site and the segmented A2 area border allowed us to locate an A2 border around the identified stereotaxic coordinates of the dye deposit. As these experiments did not have markings of the stereotaxic reference points, orientations and scaling factors of the AP and DV axes were taken as the mean of the values determined in the experiments in Figures 1–4 (orientation: 0.370 ± 0.011 radians, posterior side down; AP scaling: 3.13 ± 0.03 μm/pixel; DV scaling: 3.10 ± 0.09 μm/pixel; n = 41 mice). The genotypes of the mice included in this dataset are B6 (n = 12), CBA (n = 3), Ai9 (n = 3), PV-Cre (n = 1), and VGAT-Cre×Ai9 (n = 1).

### Calculation of relative variability in auditory cortex location

The large-scale dataset of intrinsic signal imaging mapping without stereotaxic coordinates reference points (Figure 6) includes both unpublished and published experiments from our laboratory (Aponte et al., 2021; Kline et al., 2021; Onodera and Kato, 2022). Four metrics were used to determine relative variability in the functionally-identified auditory cortical areas. 1) Cortex dimensions were measured as the total length of the combined tonotopic areas (A1, VAF, AAF, and A2) along the AP and DV axes. 2) The angle between the A1 3 kHz–AAF 3 kHz axis and the A1 3 kHz–30 kHz axis (α) and the angle between the A1 3 kHz–AAF 3 kHz axis and the A1 3 kHz–A2 3 kHz axis (β). 3) The distance between A1 and AAF borders, which is informative about the presence of a tone low-responsive center region (CTR). 4) The distribution of the area of individual regions across mice. Mice without enough trials to clearly visualize individual frequency domains were excluded from this dataset. The genotypes of the mice included in this dataset are VGAT-Cre×Ai9 (n = 86), PV-Cre×Ai9 (n = 45), CBA (n = 33), B6 (n = 30), Ai9 (n = 23), PV-Cre (n = 23), Sst-Cre×Ai9 (n = 22), Sst-Cre (n = 10), Rbp4-Cre×Ai9 (n = 9), VGAT-Cre (n = 8), Tlx3-Cre (n = 7), and Tlx-Cre×Ai9 (n = 4).

### Statistical analysis

All data are presented as mean ± SEM, except for a subset of data where mean ± SD is presented to show its variability. Statistically significant differences between conditions were determined using standard nonparametric tests in Matlab. Two-sided Wilcoxon’s rank-sum test was used for independent group comparisons. Bonferroni correction was used for multiple comparisons, and corrected *p* values were reported. Two-way or three-way analysis of variance (ANOVA) was used to examine the influence of multiple independent variables. Randomization is not relevant for this study because there were no animal treatment groups. All *n* values refer to the number of mice, except when explicitly stated that the *n* is referring to the number of area centroid pairs or mouse pairs. Sample sizes were not predetermined by statistical methods but were based on those commonly used in the field. All reported *n* are biological replications.

## Data availability

Source data are provided with this paper. Other data that support the findings of this study are available from the corresponding author upon reasonable request.

## Code availability

Custom Matlab codes used in this study will be made available from the corresponding author upon reasonable request.

## Acknowledgments

We thank Michellee Garcia and Jose Rodriguez-Romaguera for advice throughout the project and comments on the manuscript. This work was supported by NIDCD (R01DC017516), Pew Biomedical Scholarship, Whitehall Foundation, Klingenstein-Simons Fellowship (H.K.K.), Foundation of Hope (H.K.K and H.T), NINDS (F31-NS111849, T32-NS007431) (A.M.K), Toyobo Biotechnology Foundation, and Japan Society for the Promotion of Science (K.O.), and David Bray Peele Memorial Research Award from the Department of Psychology and Neuroscience, University of North Carolina at Chapel Hill (D.P.N.).

## Author Contributions

All the authors designed the project, collected data, and wrote the manuscript. D.P.N. and H.K.K. performed data analysis.

