## Supplementary Information for "Biological constraints on stereotaxic targeting of functionally-defined cortical areas"

**Supplementary Figures**

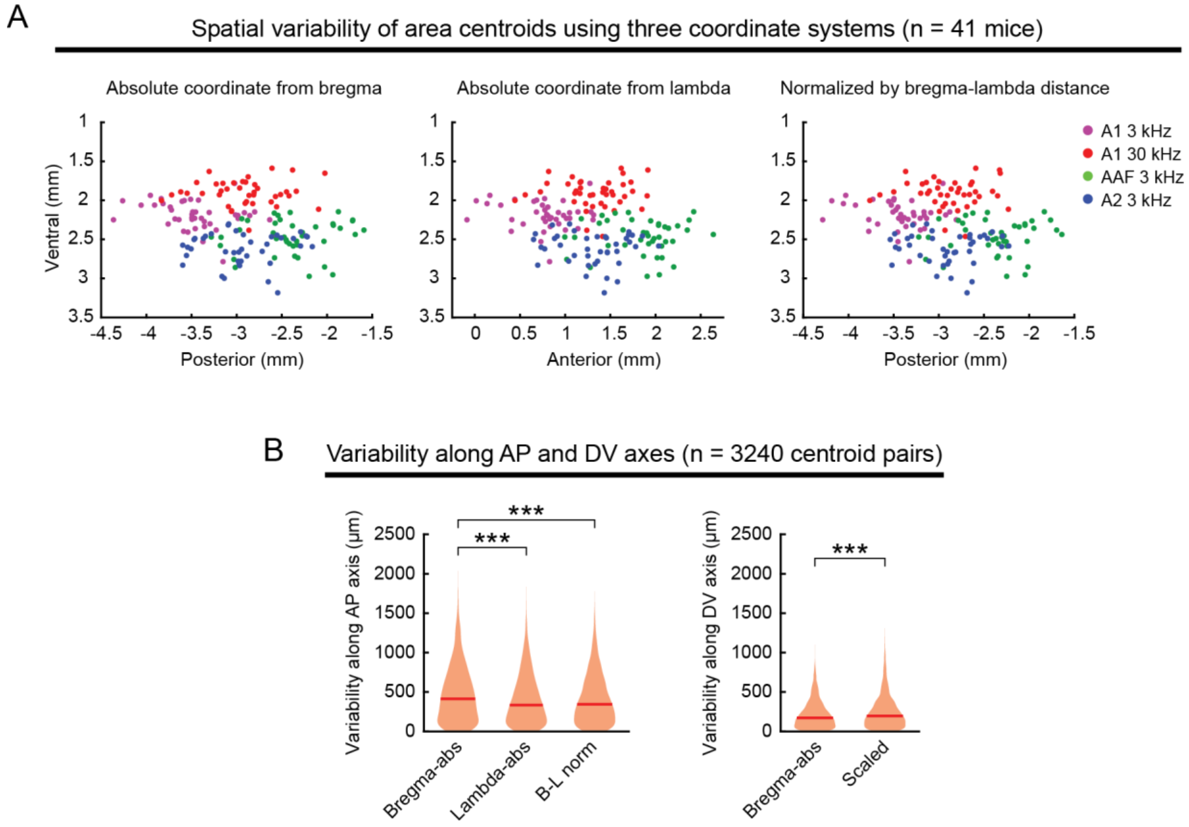


**Supplementary Figure 1. Coordinates scaled by bregma–lambda distance have the smallest variability in area centroid location. (A)** Scatter plots showing the distribution of functionally-identified frequency domain centers (A1 3 kHz, A1 30 kHz, AAF 3 kHz, and A2 3 kHz) plotted with three coordinate systems across mice (n = 41 mice). The posterior (anterior) coordinate is calculated as the absolute distance from bregma (left), the absolute distance from lambda (middle), or scaled by the distance between bregma and lambda (right, normalized to 4.2 mm). **(B)** Violin plots comparing inter-animal variability in centroid location along the anteroposterior (AP, left) and dorsoventral (DV, right) axes for each coordinate system (n = 3240 centroid pairs). For the AP axis, bregma-based absolute coordinates without size normalization resulted in larger variability than the size-normalized data. For the DV axis, scaling based on bregma–lambda distance resulted in larger variability. Red lines are median. ***$p < 0.001$. AP axis, Bregma-abs vs. Lambda-abs: $p = 6.74\times10^{-13}$, Bregma-abs vs. B–L norm: $p = 7.15\times10^{-10}$, Lambda-abs vs. B–L norm: $p = 0.814$. DV axis, Bregma-abs vs. Scaled: $p = 9.24\times10^{-8}$. Wilcoxon rank sum test. Bonferroni correction for multiple comparisons was applied for AP axis data.

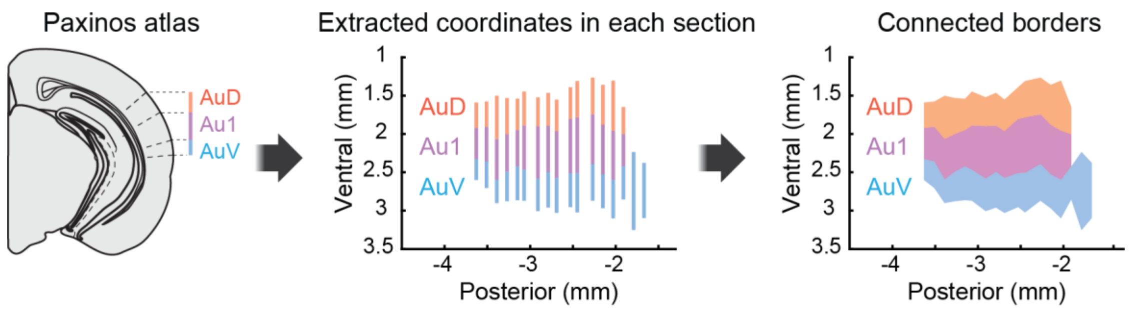


**Supplementary Figure 2. Generation of a topographical surface map of auditory cortical areas based on the Paxinos Brain Atlas.** The coordinates of the dorsal and ventral edges of Au1, AuV, and AuD were extracted from 17 atlas brain sections between −1.67 to −3.63 mm posterior from bregma. These edges were connected across sections along the AP axis.

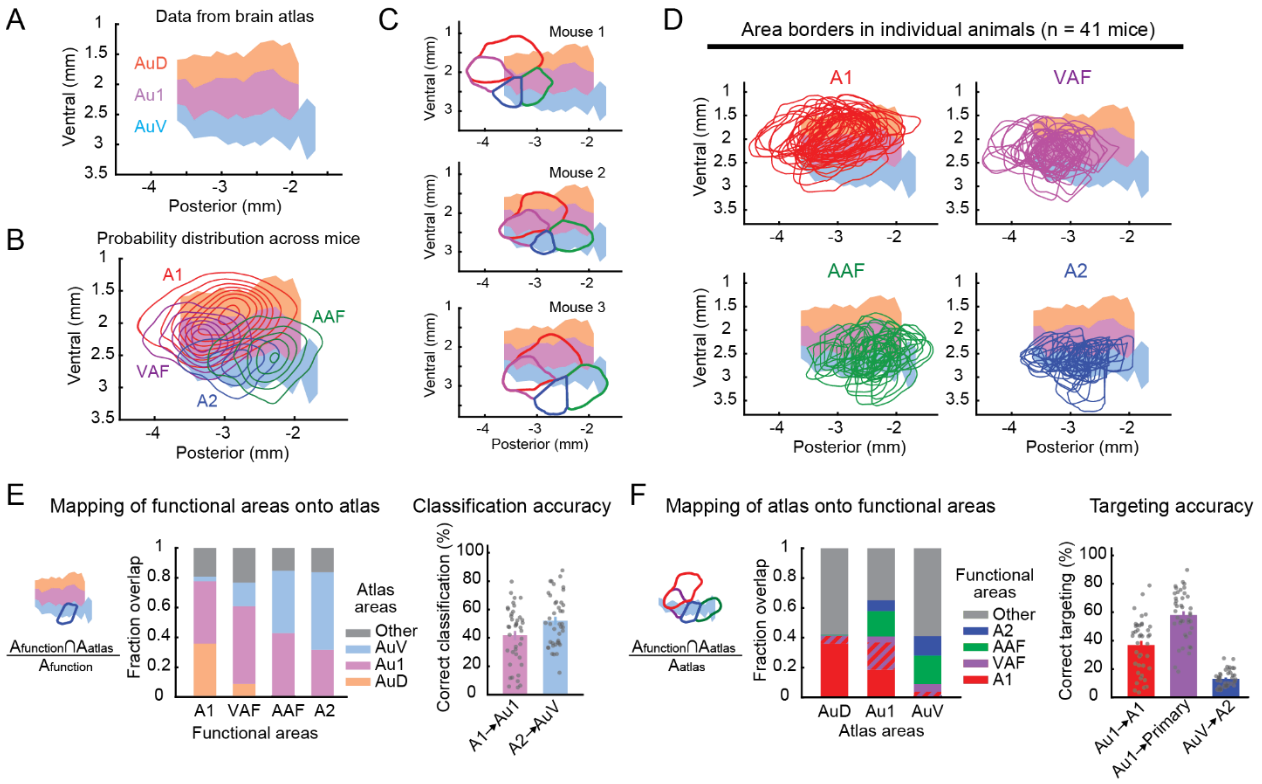


**Supplementary Figure 3. Lowering the threshold for area boundary detection to 40% peak response amplitudes does not improve targeting accuracy. (A)** A topographical surface map of auditory cortical areas based on the Paxinos Atlas. Coordinates are measured from bregma. **(B)** Probability distribution of functional auditory areas defined using a lower threshold (40% peak response amplitudes) superimposed on the atlas areas Au1, AuD, and AuV. Contours are 10% steps, starting at 10% (n = 41 mice). **(C)** Functionally-identified cortical area borders superimposed on the atlas map, showing inter-animal variability in their relationship to the atlas areas. The same three mice as Figures 1D and 2C. **(D)** Functionally-identified cortical area borders from all mice superimposed on the atlas map, shown separately for A1, VAF, AAF, and A2. **(E)** Left, fraction spatial overlap of functionally-identified areas with atlas areas. Right, classification accuracy showing the fraction of A1 contained within Au1 (A1→Au1) and A2 within AuV (A2→AuV). Each functional area tends to overlap with multiple atlas areas rather than contained within a single area, resulting in only 42 ± 2% and 52 ± 3% accuracy (n = 41 mice; mean ± SEM). **(F)** Left, fraction spatial overlap of atlas areas with functionally-identified areas. Right, targeting accuracy showing the fraction of Au1 contained within A1 (Au1→A1), Au1 within three primary areas (Au1→Primary), and AuV within A2 (AuV→A2). Using stereotaxic coordinates to target functionally-defined auditory cortex results in only 37 ± 3%, 58 ± 3%, and 13 ± 1% accuracy, respectively (n = 41 mice; mean ± SEM).

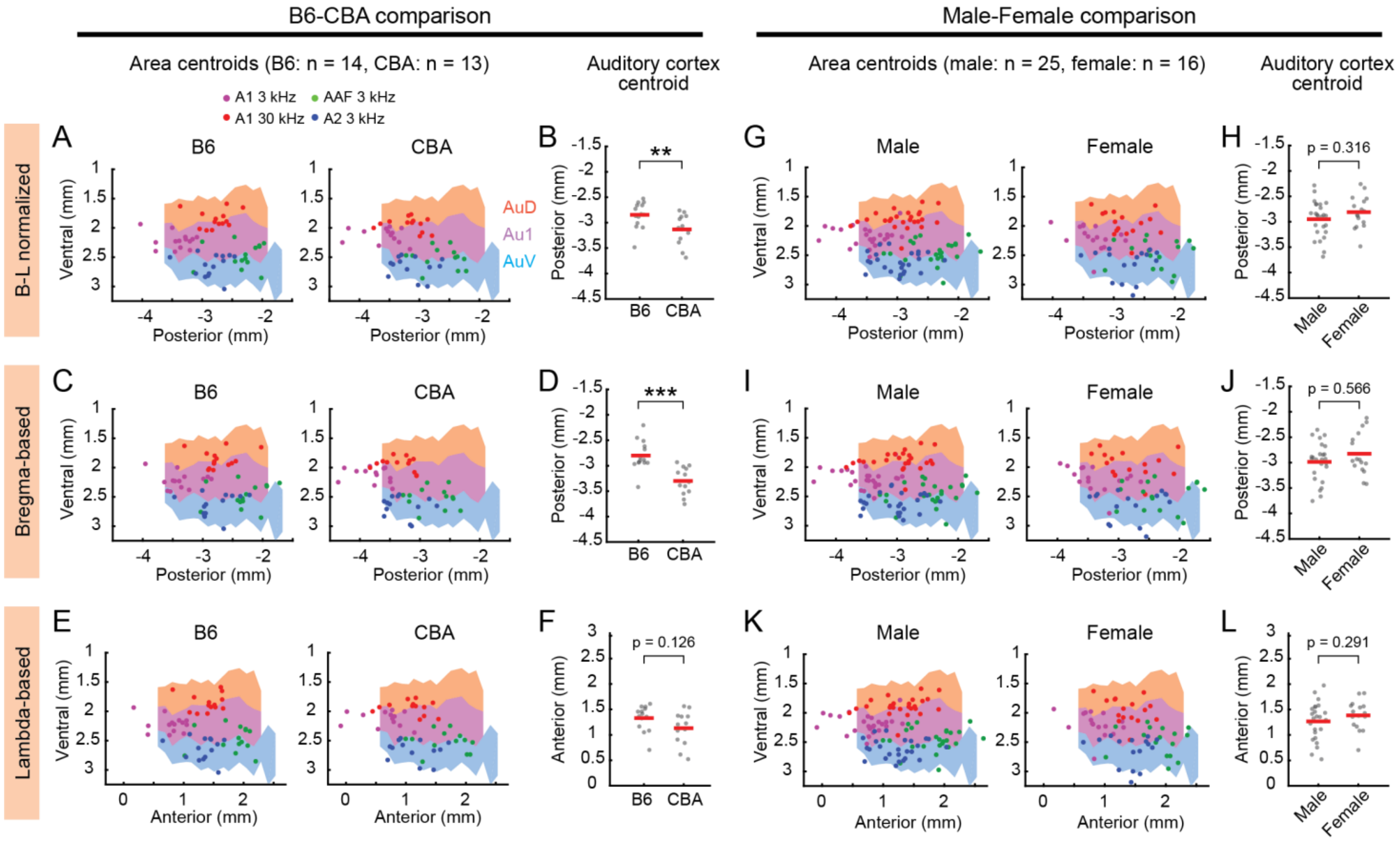


**Supplementary Figure 4. Comparison of strain- and sex-differences of functional area locations across coordinate systems. (A)** Same as Figure 3A. **(B)** Same as Figure 3B, left. Red lines are mean. **p = 0.0094, Wilcoxon rank sum test. **(C)** Distribution of functionally-identified frequency domain centroids shown separately for B6 (left, n = 14 mice) and CBA (right, 13 mice) mice, where the posterior coordinate is calculated as the absolute distance from bregma. Scatter plots are superimposed on the atlas maps. **(D)** Scatter plots showing the posterior coordinates of functionally-identified auditory cortex centroids in individual B6 and CBA mice where the posterior coordinate is calculated as the absolute distance from bregma. ***p = $1.60 \times 10^{-4}$. **(E–F)** Same as **(C–D)** but where the anterior coordinate is calculated as the absolute distance from lambda. **(G–L)** Same as **(A–F)**, but for the comparison between males (left, n = 25 mice) and females (right, n = 16 mice). This dataset includes B6, CBA, PV-Cre×Ai9, and VGAT-Cre×Ai9 strains.

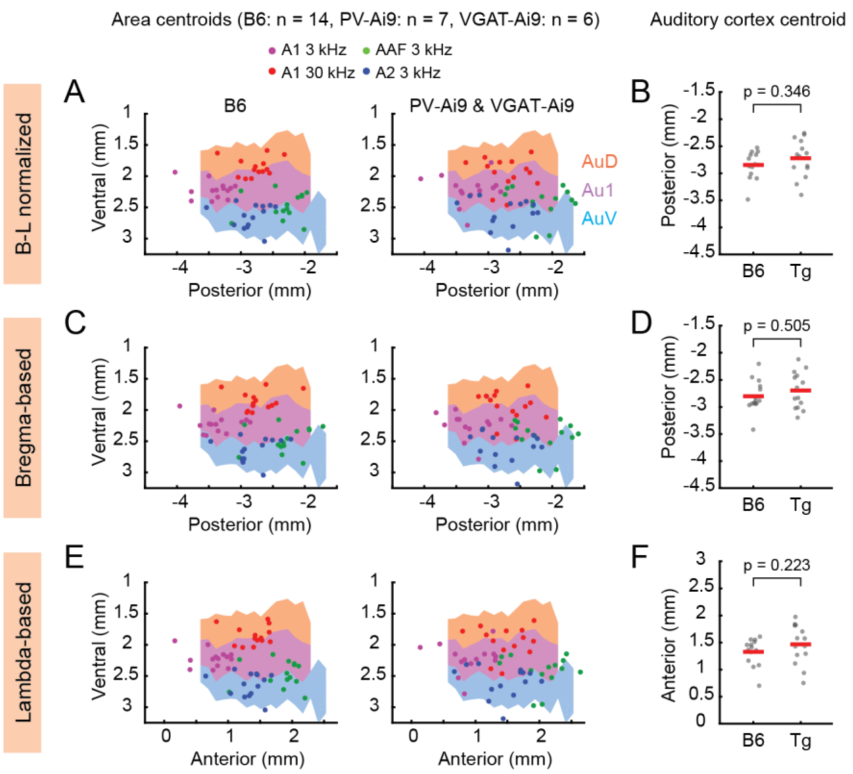


**Supplementary Figure 5. Area distributions are not significantly different between wild type B6 mice and transgenic strains. (A)** Distribution of functionally-identified frequency domain centroids shown separately for B6 mice (left, n = 14 mice) and transgenic strains (right, PV-Ai9, n = 7 mice; VGAT-Ai9, n = 6 mice). Scatter plots are superimposed on the atlas maps. The posterior coordinate is scaled to the distance between bregma and lambda. **(B)** Scatter plot showing the posterior coordinates of functionally-identified auditory cortex centroids in individual B6 and transgenic mice. Red lines are mean. Wilcoxon rank sum test. **(C–D)** Same as **(A–B),** but the posterior coordinate is calculated as the absolute distance from bregma. **(E–F)** Same as **(C–D),** but the anterior coordinate is calculated as the absolute distance from lambda.

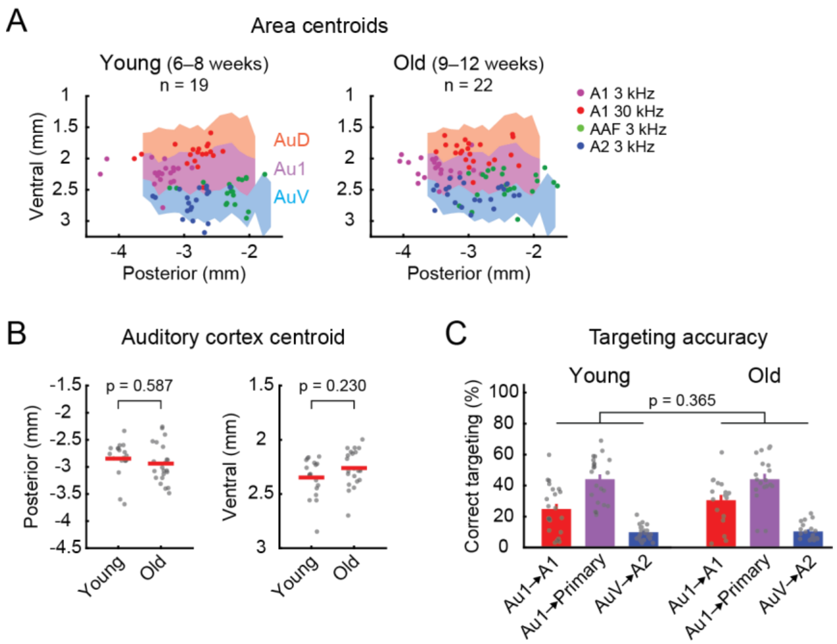


**Supplementary Figure 6. Area distributions are not significantly different between young and old mice. (A)** Distribution of functionally-identified frequency domain centroids shown separately for young (left, 6–8 weeks, n = 19 mice) and old (right, 9–12 weeks, n = 22 mice) mice. Scatter plots are superimposed on the atlas maps. **(B)** Scatter plots showing the posterior (left) and ventral (right) coordinates of functionally-identified auditory cortex centroids in individual young and old mice. Red lines are mean. Wilcoxon rank sum test. **(C)** Accuracy of using atlas-defined stereotaxic coordinates to target functionally-identified areas in young and old mice. Two-way ANOVA.

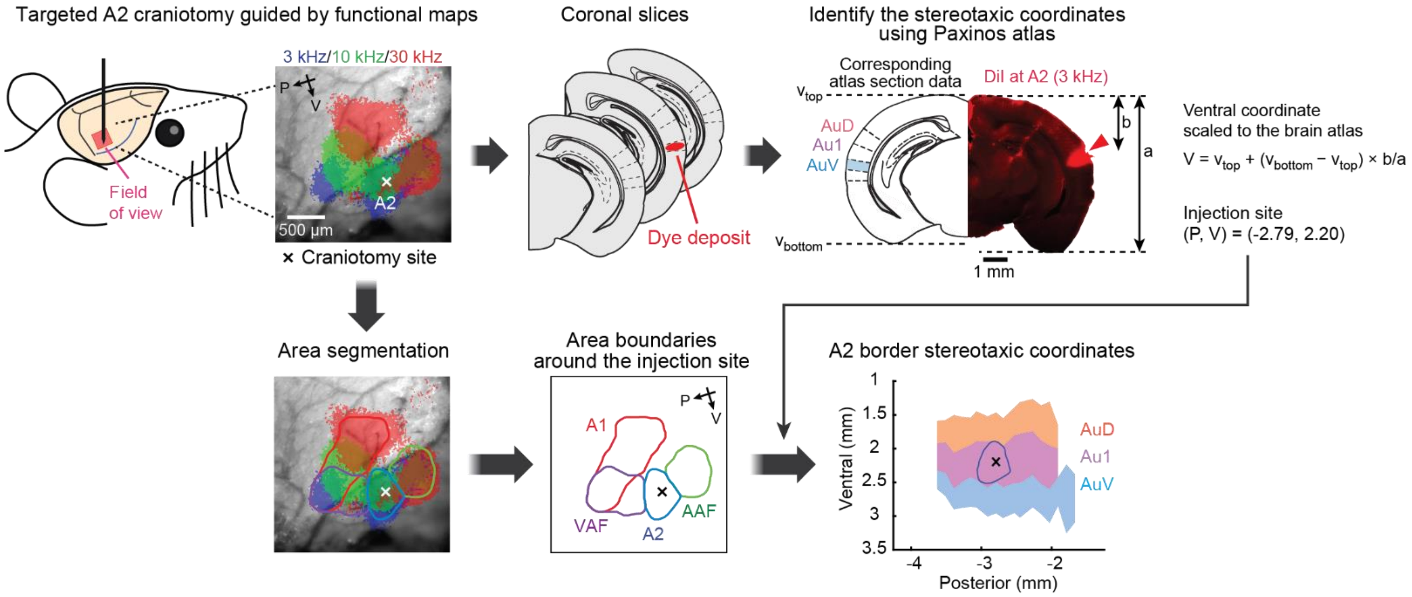


**Supplementary Figure 7. Identification of stereotaxic coordinates for dye deposits in A2.** Targeted manipulations were performed in A2 (see Methods for additional detail) guided by intrinsic signal imaging. After sectioning the brains, the section with the strongest fluorescence was identified as the center of the marking and used to identify stereotaxic coordinates with the Paxinos Brain Atlas. The posterior coordinate of the dye deposit was determined by identifying the atlas section with the corresponding morphology. The ventral coordinate was determined by vertically scaling the histology section to the height of the atlas section. The craniotomy site (shown as a cross) within the intrinsic signal imaging map was recorded during the experiment and used to draw an A2 border around the identified stereotaxic coordinates of the dye deposit. Finally, this functional area boundary was superimposed on the atlas map.

**Supplementary Protocol**

**Intrinsic Signal Imaging of Cortical Sensory Responses Through Intact Skull in Mice**

**Materials and Reagents**

1. 70% ethanol (any vendor)
2. 10% Povidone-iodine (Betadine; any vendor)
3. Isoflurane (any vendor)
4. Ocular lubricant (any vendor)
5. Sterile phosphate-buffered saline, pH 7.2 (any vendor)
6. Razorblade for shaving (any vendor)
7. Sterile cotton tip applicators (any vendor)
8. Chlorprothixene, 1.5 mg/kg body weight (Sigma-Aldrich)
9. Silicone sealant (KWIK-CAST (WPI) or Body Double-Fast Set (Smooth-On))
10. Cyanoacrylate tissue adhesive (Vetbond (3M))
11. Dental cement (Jet Denture Repair Powder and Liquid (Lang) for temporary adhesion; Super-Bond C&B (Sun Medical) for chronic implantation)
12. Analgesics (Meloxicam 5 mg/kg body weight, or as specified by individual institutions)
13. Antibiotics (Enrofloxacin 10 mg/kg body weight, or as specified by individual institutions)
14. Anti-inflammatory drugs (Dexamethasone 2 mg/kg body weight, or as specified by individual institutions)

**Equipment**

1. Stereotaxic frame (Kopf Instruments Model 1900)

2. Dissecting microscope (Leica)
3. Feedback-controlled temperature controller (FHC)
4. Isoflurane vaporizer (VetEquip V-1 Table-top lab animal anesthesia system)
5. Hot beads sterilizer (FST)
6. Dumont #5 forceps (FST)
7. Fine scissors, straight, 9 cm (FST)
8. Scalpel handle #3 (FST)
9. Scalpel blade #11 (any vendor)
10. Metal alligator clips (any vendor)
11. Microscope cover glass (Fisher Scientific, 22×22-1)
12. Diamond scribe (Fiber Instrument Sales)
13. Custom stainless-steel head bar (3×19×1.2 mm) and clamps, or any head-fixation system of choice
14. Custom imaging stage to hold head-fixed mouse
15. Sound isolation chamber (Gretch-Ken Industries)
16. Free-field electrostatic speaker system (Tucker-Davis Technologies)
17. Custom tandem-lens macroscope (composed of Nikkor 35mm 1:1.4 and 135mm 1:2.8 lenses) with four-axis manipulator.
18. 530 nm LED (Thorlabs M530F2)
19. 625 nm LED (Thorlabs M625F2)
20. 12-bit CMOS camera (Dalsa DS-1A-01M30) and associated image acquisition computer
21. Bpod (Sanworks) and associated sound stimulus generation computer
22. Matlab (Mathworks)
23. ImageJ (Fiji)
24. ImageJ plugin, IO and VSD Signal Processor (https://murphylab.med.ubc.ca/io-and-vsd-signal-processor/) (Harrison et al., 2009)

### Procedures

Section A describes surgical procedures for intrinsic signal imaging (Aponte et al., 2021; Kato et al., 2017, 2015; Kline et al., 2021) with temporary head bar implantation, which is followed by re-closure of the scalp. Alternatively, the head bar may be chronically attached to the skull during these procedures to form a cement head cap instead of putting the scalp back. For example, chronic head bar implantation is suitable for performing targeted *in vivo* electrophysiological recording following functional mapping. In that case, at least one day of recovery period is recommended due to the long-lasting sedative effect of chlorprothixene.

#### A. Surgical procedures with temporary head bar implantation

1. Anesthetize the mouse in the induction chamber with isoflurane (4%) vaporized in oxygen (1 L/min). After the mouse has reached deep anesthesia (~1 Hz breathing), weigh the mouse and move it to the surgery station.
2. Fix the mouse with the nose cone and keep its body temperature at 34–36 ºC on a feedback-controlled heating pad. Isoflurane (1.2–2% in oxygen; gradually ramp down during the surgery by monitoring the respiration rate) is delivered through the nose cone. Confirm anesthetic depth by testing the toe-pinch reflex.

   *NOTE: It is critical to keep the isoflurane level at the minimum required level. Increased anesthesia depth critically reduces intrinsic signals.*
3. Cover the eyes with ocular lubricant to protect them from drying out (Figure 1A). Push down the whiskers with lubricant to keep them away from the surgical area.
4. Shave the top and right side of the mouse head and disinfect the area by applying ethanol and Povidone-iodine (Figure 1B).
5. Make an incision along the midline and expose the skull by holding the right scalp with an alligator clip (Figure 1C).

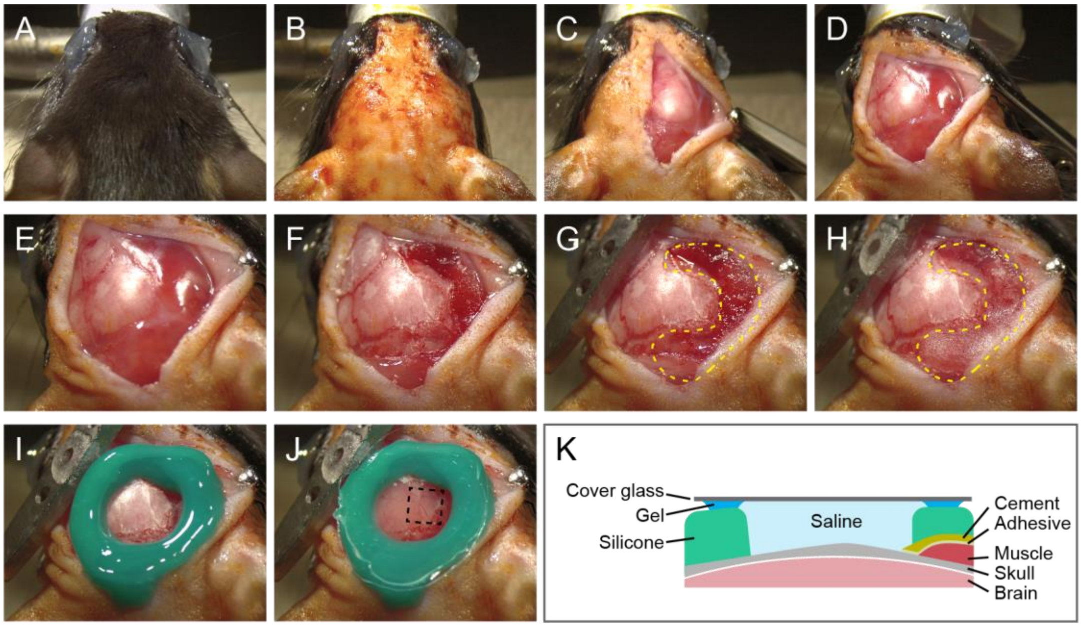


**Figure 1. Surgical procedures for intrinsic signal imaging of the auditory cortex.**
**(A–C)** Dorsal views of the mouse head before surgery (A), after shaving (B), and after skull exposure (C). **(D)** Lateral view after rotating the head. **(E, F)** Lateral views of the temporal region before (E) and after (F) removing the muscle overlying the auditory cortex. **(G, H)** Lateral views after covering the wound (yellow dotted lines) with tissue adhesive (G) and dental cement (H). A stainless-steel head bar is cemented to the skull. **(I, J)** Views around the auditory cortex after putting silicone sealant (I) and a glass window (J). A dotted square in (J) indicates the field of view of imaging. **(K)** Schematic coronal view of the implanted window.

6. Rotate the head and remove the temporal muscle overlying the region of interest (ROI) around the right auditory cortex (Figure 1E, F). Use sterile cotton tip applicators to wipe away fluid and remove connective tissue around the ROI by gently scratching with a scalpel.

   *NOTE: The purpose of scratching here is to remove a thin layer of tissue and not to make grooves. Excessive scratching will cause bleeding and obscure intrinsic signals.*

7. Seal the wound with tissue adhesive to prevent bleeding and attach a stainless-steel head bar (or any head-fixation apparatus of choice) to the skull (Figure 1G).

8. Cover the area around the ROI (but not the ROI) with a thin layer of dental cement (Figure 1H). Put a small amount of dental cement to secure the head bar on the skull. Alternatively, if chronic head bar implantation is desirable, cover the entire top surface of the skull with dental cement.

   *NOTE: The adhesive and the cement prevent the diffusion of blood, which introduces significant noise to the signal, into the saline during imaging.*

9. Make a well surrounding the ROI with silicone sealant (Figure 1I).
10. Prepare a round glass window by cutting cover glass with a diamond scribe.
11. Put petroleum gel on the silicone well while avoiding direct contact with the skull, apply degassed PBS into the well, and seal with a glass window (Figure 1J, K).

    *NOTE: Saturation with PBS keeps the skull transparent during imaging. Tight sealing with petroleum gel and a glass window is critical to prevent evaporation. Instead of saturation with PBS, forming a thin and smooth layer of transparent cement over the ROI also allows imaging, depending on the experiment. Degassing of PBS helps reduce air bubbles that appear as the solution warms up.*
12. Run intrinsic signal imaging and analyze the data (Figure 2). See Sections B and C below for details. Imaging and analysis take less than 30 min and 10 min, respectively.
13. After intrinsic signal imaging, remove the glass window, petroleum gel, and silicone sealant, and clean the skull surface.
14. Conduct targeted experimental procedures, such as craniotomy and virus injection, as necessary (Figure 3Ai-ii and 3B). Depending on the experimental goal, a similar procedure can be used to target

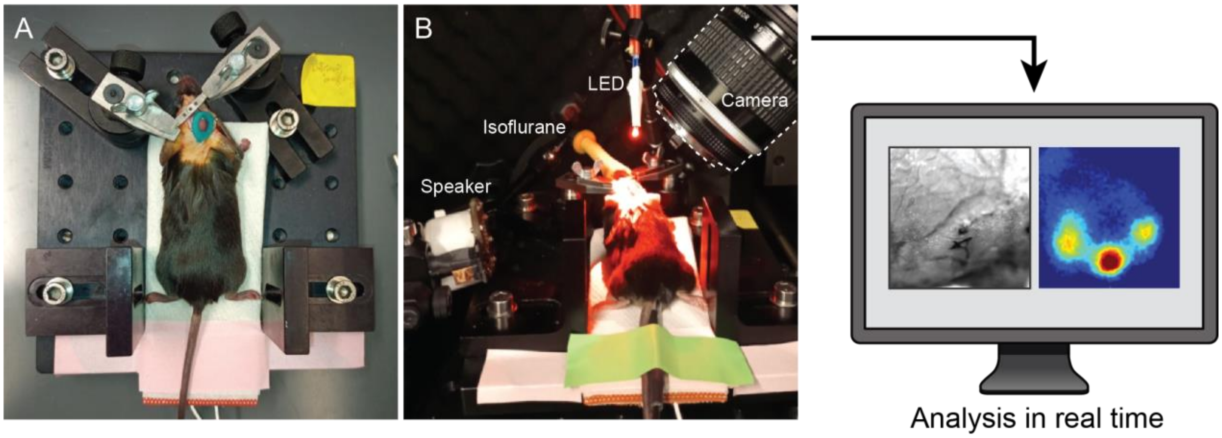


**Figure 2. Image acquisition setup. (A)** Top view of a mouse head-fixed on an imaging stage with a feedback-controlled heating pad. **(B)** Left, intrinsic signal imaging in a sound isolation chamber. The signals are imaged with a tandem-lens macroscope using a red LED illumination. Right, image acquisition and simple mapping analyses can be performed in real-time (see Figure 4A).

electrophysiological recordings (Figure 3C) and two-photon calcium imaging (Figure 3D).

15. Remove the clip and gently remove the tissue adhesive and dental cement. Remove the head bar with a scalpel blade (Figure 3Aiii).
16. Put back the scalp and close the wound with tissue adhesive (Figure 3Aiv).
17. Inject analgesics, antibiotics, and anti-inflammatory drugs subcutaneously, as determined by individual institutions.
18. Recover the mouse on a heating pad and monitor for its recovery as specified by the institution. Chlorprothixene has long-lasting sedative effect, which could remain up to several hours.

    *NOTE: The amplitudes of intrinsic signals critically depend on the anesthesia depth and the duration of the surgery. Ideally, the entire surgery, imaging, and analysis should be completed within 1.5–2 hours to obtain clear signals.*

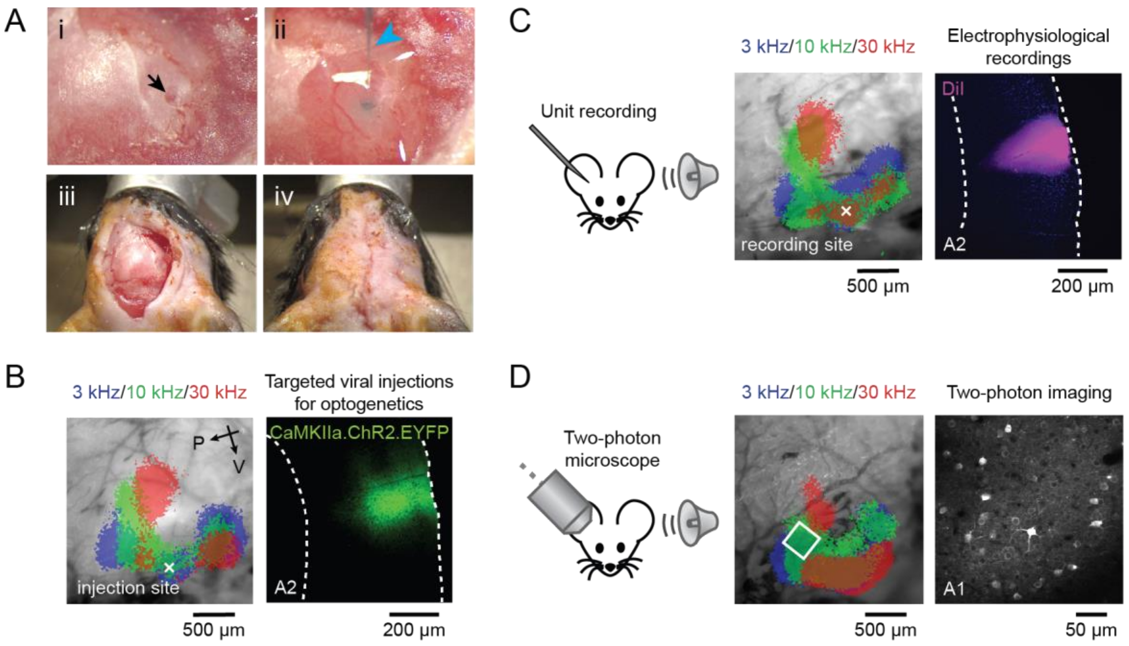


**Figure 3. Post-imaging application options.** **(A)** Optional area-targeted craniotomy (i) and virus injection (ii). (iii) After removal of the cement and head bar. (iv) After the closure of the scalp using tissue adhesive. Black arrow: craniotomy. Cyan arrowhead: injection glass pipette. **(B)** Example targeting of viral injection to A2. Optogenetic tool channelrhodopsin is expressed via AAV.CaMKIIa.ChR2.EYFP. White cross: craniotomy site. **(C)** Example targeting of unit recording to A2. The recording location is identified in post-recording histology by DiI on the probe. **(D)** Example targeting of two-photon calcium imaging to A1 mid-frequency domain. White square: two-photon imaging field of view.

### B. Intrinsic signal imaging

1. Inject the mouse with chlorprothixene (1.5 mg/kg body weight) subcutaneously prior to imaging. Chlorprothixene helps reduce the isoflurane concentration and minimizes isoflurane's negative influence on hearing (Ruebhausen et al., 2012).
2. Head-fix the mouse on a stage using any head fixation apparatus of choice (Figure 2A).
3. Place the stage in a sound isolation chamber (Figure 2B). Deliver low-concentration isoflurane (0.8%) through a mask and keep the body temperature at 34–36 ºC using a feedback-controlled heating pad.
4. Focus on the skull surface around the ROI with a tandem-lens macroscope (Ratzlaff and Grinvald, 1991) and a 12-bit CMOS camera. We image 2.3 × 2.3 $mm^2$ area in 717 × 717 pixels at a 16 Hz sample rate. Lower sample rates are acceptable as the intrinsic signal has slow kinetics.

   *NOTE: 12-bit or higher bit depth is necessary to visualize subtle changes in the reflectance.*
5. Acquire a surface vasculature image using green (530 nm) illumination.
6. Lower the objective 400–500 µm along the optical axis to focus on L4–L5.

   *NOTE: Focusing on superficial layers results in weaker sound-evoked signals and larger contamination from blood vessel artifacts.*
7. Switch to red (625 nm) illumination and set it to the highest intensity just below saturation of the camera, as intrinsic signal imaging detects a reduction in reflectance.
8. Acquire videos while presenting sound stimuli. Each trial consists of a 1-s baseline followed by a 1-s tone stimulus (75 dB SPL pure tone with a frequency of 3, 10, or 30 kHz) and a 30-s inter-trial interval. If surgery is successful, coarse localization of signals to tonotopic areas is visible in single trials. However, averaging the results over 5–20 trials improves the signal-to-noise ratio.

   *NOTE: Inter-trial interval needs to be at least 20 seconds to avoid contamination between trials.*

### C. Data analysis

1. Calculate response maps in individual trials as $(R_{tone}-R_0)/R_0$, where $R_{tone}$ and $R_0$ are the average image during the response period (0.5–2 s from the tone onset) and the baseline, respectively (Figure 4A).
   *NOTE: ImageJ plugin, IO and VSD Signal Processor can be used (https://murphylab.med.ubc.ca/io-and-vsd-signal-processor/) (Harrison et al., 2009).*
2. Average the response maps across trials for each sound. Additionally, deblurring can be applied with a 2-D Gaussian window (σ = 200 μm) using the Lucy-Richardson deconvolution method (Issa et al., 2014; Romero et al., 2020).
3. For visualization, the signals can be binarized and overlaid across sounds (Figure 4B top).
4. For unbiased area segmentation, perform semiautomated identification of frequency domain boundaries for A1, AAF, VAF, and A2 (Figure 4B bottom).
5. Superimpose the resulting signal maps on the surface vasculature image.

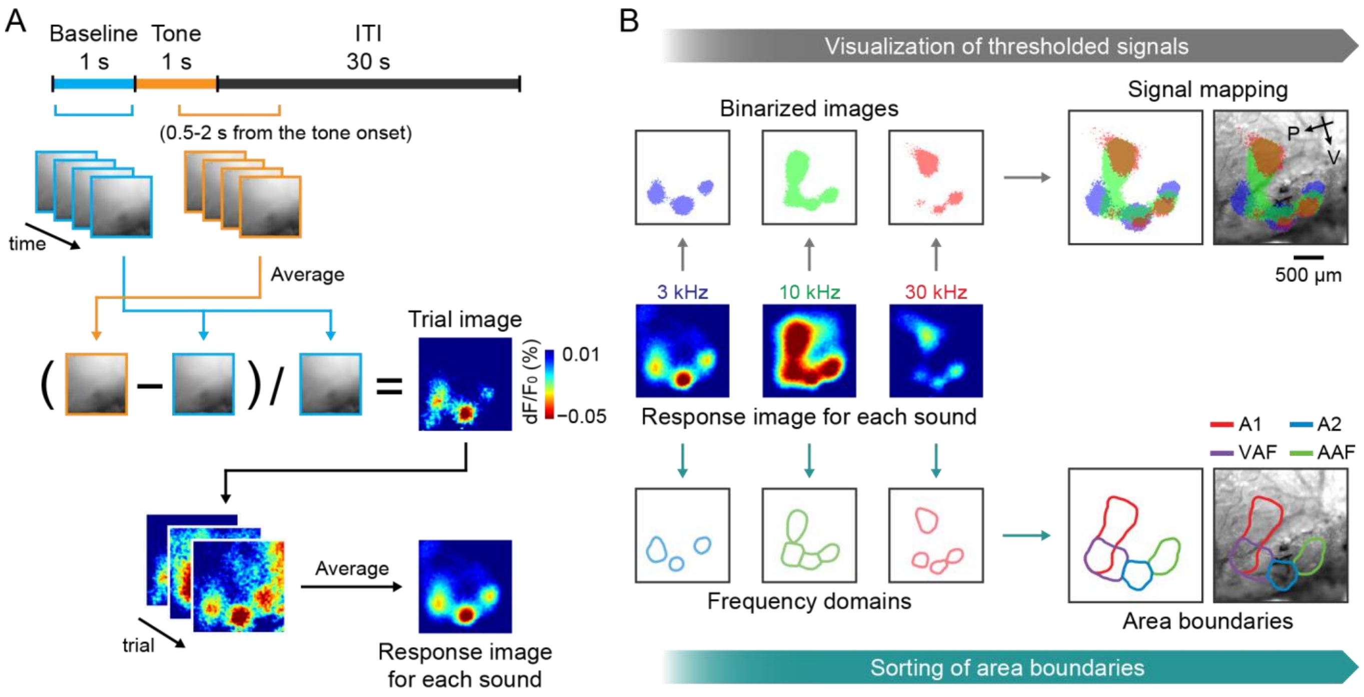


**Figure 4. Image processing for intrinsic signals of tone-evoked responses.**
**(A)** Schematics showing the trial structure and data analysis workflow. **(B)** Schematics showing the options for visualization of intrinsic signal maps. Top, visualization of thresholded response maps for individual sounds. Bottom, semiautomated determination of functional area boundaries. Maps are overlaid onto the surface vasculature images.
